# A novel minimally invasive and reproducible large animal ischaemia-reperfusion-infarction model: methodology and model validation

**DOI:** 10.1101/2023.03.02.530817

**Authors:** Charlene Pius, Barbara Niort, Emma J. Radcliffe, Andrew W. Trafford

**Affiliations:** Division of Cardiovascular Science, School of Medical Science, Faculty of Biology Medicine and Health, University of Manchester, Manchester Academic Health Science Centre, United Kingdom

## Abstract

Ischaemic heart disease remains a leading cause of premature mortality and morbidity. Understanding the associated pathophysiological mechanisms of cardiac dysfunction arising from ischaemic heart disease and the identification of sites of novel therapeutic intervention requires a preclinical model that reproduces the key clinical characteristics of myocardial ischaemia, reperfusion and infarction. Here we describe and validate a refined and minimally invasive translationally relevant approach to induce ischaemia, reperfusion and infarction in the sheep. The protocol uses clinical cardiology devices and approaches and would be readily adopted by researchers with access to standard fluoroscopic instrumentation. In addition to being minimally invasive, the major refinements associated with the described methodology are the implantation of an intracardiac defibrillator prior to coronary engagement and use of an antiarrhythmic medication protocol during the procedure. These refinements lead to a reduction of intraoperative mortality to 6.7 %. The model produces key characteristics associated with the 4^th^ Universal Definition of Myocardial Infarction including electrocardiographic changes, elevated cardiac biomarkers and cardiac wall motility defects. In conclusion, the model closely replicates the clinical paradigm of myocardial ischaemia, reperfusion and infarction in a translationally relevant large-animal setting and the applied refinements reduce the incidence of intraoperative mortality typically associated with preclinical myocardial infarction models.

## Introduction

Cardiovascular diseases, mainly ischaemic heart diseases (IHD), are the leading cause of mortality in the United Kingdom^1, 2^ and worldwide^3^. Coronary artery disease (CAD) is one of the most important causes of IHD^4^ and is characterised by a reduction in the volume of perfusion to the heart (i.e., ischaemia) or even its complete cessation (i.e., infarction)^5–7^. Acute myocardial infarction (MI) is defined as myocardial necrosis in the context of myocardial ischaemia, which can be transmural, involving all three layers of the heart (endocardium, mid-myocardium, and epicardium) or non-transmural, typically sparing the epicardium^5–11^. Classically, non-transmural MI present clinically with non-ST segment elevation (NSTEMI) on the electrocardiogram (ECG) are treated primarily with anti-platelet treatment, peri-interventional anticoagulant treatment, and coronary angiography with a view to revascularization within 72h^11^. Transmural MI are associated with ST segment elevation (STEMI) on the ECG and are treated with primary percutaneous coronary intervention (PPCI)^12^. Complications following MI include ventricular arrhythmias (VA)^13, 14^ and heart failure (HF)^15^, which can manifest early or late^15–17^. VA can occur in the form of ventricular tachycardia (VT)^14, 18^ or ventricular fibrillation (VF). These life-threatening complications necessitate increased research to identify novel therapeutic targets that may ultimately alter prognosis following MI and ultimately this necessitates translationally relevant animal models.

When studying a disease, especially the precise molecular aspects of dysregulation, the animal model should ideally be as clinically relevant as possible. Occluding a coronary artery in an experimental animal can induce a reaction comparable to an acute MI caused by atherosclerosis or thromboembolic events. Most of the current small (mouse or rat)^19^ and large (pig, sheep or dog) mammal models of cardiac dysfunction from ischaemia consist of the permanent ligation of the LAD^20–24^ (for review see^25, 26^). Whilst they are reliable models for inducing tissue damage and HF, they do not accurately reflect the clinical setting occurring as a result of a reperfusion of the occluded vessel during PPCI. Other models use intracoronary injections of thrombogenic material causing a permanent occlusion and therefore, in the same way, do not reflect the typical clinical situation. Importantly, the reperfusion phase may also be associated with cardiac dysfunction known as ischaemia-reperfusion injury (IRI) including arrhythmias, myocardial stunning and vascular obstruction^27^. None of these potential sequelae occur in models of total and permanent occlusion.

The development of percutaneous transluminal coronary angioplasty balloon catheters to treat coronary atherosclerotic stenosis has led to the creation of less invasive ischaemia-reperfusion animal models. However, all these models still present with a low survival rate^20, 28, 29^, predominantly owing to arrhythmic death or low cardiac output state^28^. This high mortality rate is at odds with key elements of the 3R’s principles of animal research^30–32^ (Reduction; Refinement and Replacement), making the establishment of a large mammal MI model more challenging.

The heart of large animals shares many electrophysiological and contractility similarities to humans, which is why modelling cardiac diseases in these species generally better reflects human pathologies and thus drug and interventional effects than in small animals^33^. Additionally, large animal models also allow for the implementation of minimally invasive approaches and use of clinical grade materials and devices thus conferring an inherent refinement aspect of 3R’s considerations. Canine and porcine models using balloon occlusion of coronary arteries already exist. However, both models present limitations. Studies suggest that the vascular architecture of the porcine and canine heart differs from that of a human heart, and hence may be less representative of the remodelling that occurs in a human with ischaemic heart disease. Specifically, when compared to human, pig and sheep, rodent and canine hearts are known to have a greater collateral network which can affect the ability to form a predictable infarct size^34–36^. Furthermore, pig models are associated with a high rate of sudden death caused by ventricular arrhythmias following MI (reviewed in^37^). Lastly, an additional logistical consideration is that husbandry can become very difficult as pigs grow rapidly to large size. The ovine heart is one of the closest to the human, and therefore is accepted as a good pre-clinical animal model for cardiovascular research^38^. A clear scientific protocol for the experimental induction of ischaemia/infarction-reperfusion injury and study of STEMI in sheep is lacking. It is essential to develop comprehensive, reproducible, reliable protocols and criteria for knowledge and skill transfer, as well as to ensure that investigations can be replicated with minimal animal suffering to ensure good proximity to the clinical scenarios the in vivo modelling is attempting to reproduce.

Here, we have developed a minimally invasive large mammal ischaemia/infarction-reperfusion model in sheep, which is more clinically relevant, with a considerably lower mortality rate (6.7%) than other previously reported large animal models. To that end, we refined a prophylactic intra-operative anti-arrhythmic drug protocol and added a surgical step involving the implantation an internal cardiac defibrillator to assist with rapid defibrillation of ventricular arrhythmias.

## Results

### Defining a model of infarction-reperfusion large mammal

A total of 28 female Welsh Mountain sheep aged ∼18 ± 6 months weighing 38 ±1.2 kg were used in this study. Following carotid cannulation, a 6F JR4 guide catheter was advanced down the 6F haemostatic sheath to reliably engage the left coronary ostium. After left coronary artery angiography, the balloon was successfully placed below the second diagonal branch of the LAD (supplementary figure 1). Only 2 of the 28/30 sheep died following MI induction from intractable ventricular fibrillation. Total mortality was 6.7 % all occurring intraoperatively before reperfusion.

Using a minimally invasive coronary angiographic technique, the infarct was created by inflating an intra-coronary balloon for 90 minutes to occlude blood flow followed by reperfusion. Both occlusion and reperfusion were confirmed by angiography. In order to validate the model, we set out to achieve the key criteria set out in the 4^th^ universal definition where MI is defined as myocardial necrosis in the context of myocardial ischaemia resulting in an elevation of a cardiac biomarker such as cardiac troponin I (cTnI)^39^ with at least one value above the 99^th^ percentile upper reference limit. The diagnosis of MI requires additional criteria, including at least one of the following: (1) clinical symptoms of ischaemia including chest discomfort; (2) electrocardiographic (ECG) ischaemic features such as the development of new changes in the ST and T segment (ST-T) or new left bundle branch block (LBBB); (3) new pathological Q waves; (4) imaging evidence of reduced cardiac wall contractility; and/or (5) evidence of an intracoronary thrombus visualised via angiography or on autopsy^40, 41^. Specific objective assessment of chest discomfort is clearly impossible to achieve in animal models, however each of the remaining four criteria are assessable with readily applicable methods or investigations. We will, in turn, consider each of the assessable 4^th^ Universal Definition criteria achieved using the minimally invasive model of STEMI presented here.

### Temporal changes in serum cTnI

In this model, cTnI serum levels were easily and rapidly evaluated^42^ using whole blood samples and a point-of-care device (see Methods; upper limit of detection 50 ng/ml). Measurements were made at various stages during the procedure (Fig 1). At baseline, all animals had near-zero values. Following the inflation of the angioplasty balloon, we observed an elevation in a cTnI (p < 0.0001), displaying the typical rise and fall pattern seen in STEMI patients^43–46^. The cTnI levels rose significantly during reperfusion (1.2 ± 1ng/ml; p < 0.01) and continued to rise at 30 minutes post-reperfusion (27.4 ± 3 ng/ml; p < 0.0001) with a peak observed at a measurement time of 90 minutes post-reperfusion (46 ± 1.8 ng/ml; p < 0.0001) when compared to baseline (0.03 ± 0 ng/ml). The levels started to decline by day 2 to 4 (21.5 ± 3 ng/ml; p < 0.0001 vs baseline) with a further decline after 1 week (0.6 ± 0.2ng/ml; p < 0.01 vs baseline) and returning to baseline values by week 3 (0.03 ± 0.0 ng/ml) and remaining low until the end of the study (week 8; 0.01 ± 0.0 ng/ml) (Fig. 1).

**Fig. 1.**
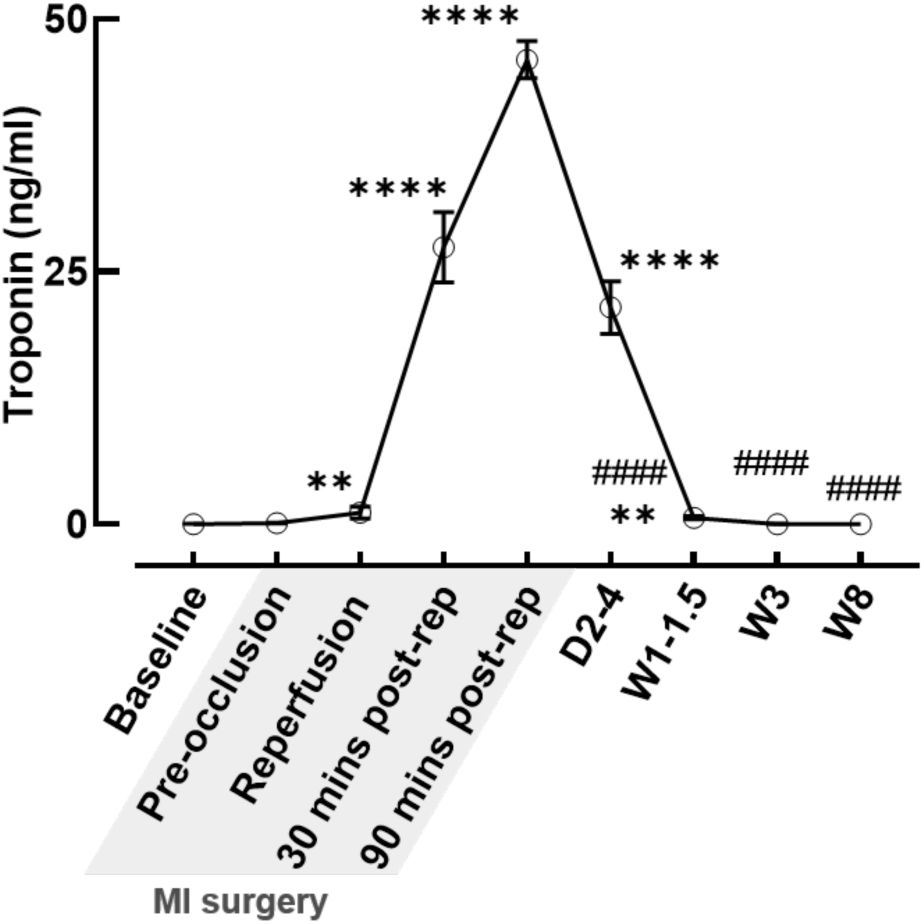
cTnI measurements. cTnI measurements at baseline, intraoperatively and on day 2 - 4, 1 - 1.5 weeks, 3 weeks and 8 weeks demonstrating the expected rise and fall pattern seen in MI. *N*, baseline= 28; pre-occlusion = 28; immediate reperfusion = 27; at 30 mins post-reperfusion = 28; 90 mins post-reperfusion = 28; Days 2 - 4= 28; Weeks 1 - 1.5= 20; Week 3 = 9; Week 8 = 10. **** p < 0.0001 ** p < 0.01 vs baseline; #### p < 0.0001 vs 90 minutes post-reperfusion by Kruskal-Wallis test.

### Electrocardiographic changes following myocardial ischaemia-reperfusion injury

The intention with this model was to create an ST segment elevation MI thus most closely resembling the clinical cohort of patients being treated with PPCI. The ST segments started to rise upon coronary occlusion (0.07 ± 0.02 mV) with a statistically significant increase seen on reperfusion (0.13 ± 0.03 mV; p < 0.0001) compared to baseline (0.02 ± 0.00 mV). ST segment normalisation was already evident at the end of surgery (0.05 ± 0.01 mV; p < 0.05 vs reperfusion) (Fig. 2a and b). The dynamic pattern of these ST segments supports the diagnosis of MI^41, 47^. Intraoperative ventricular arrhythmias requiring intracardiac defibrillation were observed in 8 of 27 animals.

**Fig. 2.**
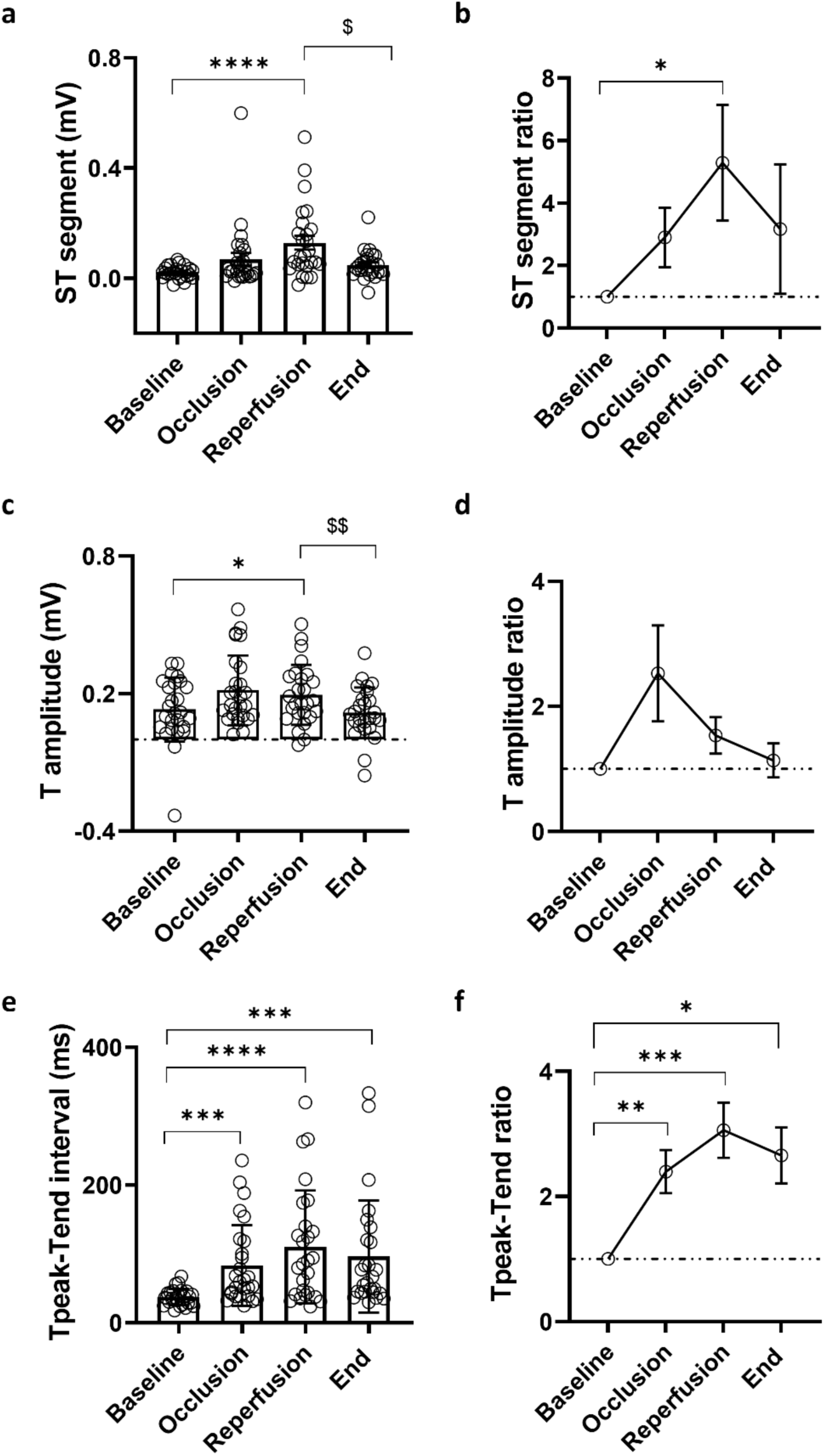
Electrocardiographic changes following ischaemia-reperfusion injury. ST segment changes as **a**, absolute measurements and **b**, difference in height compared to baseline. T wave amplitude changes as **c**, absolute measurements and **d**, ratio compared to baseline measurements. Tpeak-Tend interval changes as **e**, absolute values and **f**, ratio values compared to baseline. *N*, baseline = 25 - 27, occlusion = 25 - 27, reperfusion = 25 - 26, end = 25 -26. **** p < 0.0001, *** p < 0.001, ** p < 0.01, * p < 0.05 vs baseline, $$ p < 0.01, $ p < 0.05 vs reperfusion by mixed effects model analysis and RM one-way ANOVA.

The T wave amplitude peaked on reperfusion (0.19 ± 0.03 mV; p < 0.05) compared to baseline (0.13 ± 0.03 mV) with a gradual return to near normal values by the end of surgery (0.12 ± 0.02 mV; p < 0.01 vs reperfusion) (Fig. 2c and d). T peak amplitude increment could be a consequence of interstitial hyperkalaemia from myocardial ischaemia^42, 43^.

We also took the opportunity to calculate the T-peak Tend interval, which is suggested as a measure of transmural dispersion of repolarization^40, 41^. Upon coronary occlusion, the Tpeak-Tend interval more than doubled (83 ± 11 ms vs 37 ± 2.3ms ; p < 0.001) with maximal prolongation seen on reperfusion (110 ± 16ms; p < 0.0001) declining by the end of surgery (96 ± 16 ms; p < 0.001) whilst remaining prolonged compared to baseline (Fig. 2d and e).

### Change in cardiac contractile function and planimetry

Left ventricular ejection fraction (EF) was estimated at intervals as described in the methods section. Pre-surgery EF was 73 ± 1.4 % and declined following MI being 58 ± 2.1% at 1 – 1.5 weeks (p < 0.0001), 60 ± 3.1% (p < 0.05) and 52 ± 1.7 % at 8 weeks post MI (p < 0.0001) demonstrating a 21% decline in EF compared to baseline (p < 0.0001) (Fig. 3a). While the final EF value at 8 weeks was higher than those mentioned in other studies^44^, the total reduction in EF of 21% was bigger than the 14% described in a previous study^45^. We also measured fractional shortening (FS), as a measure of contractility and this parameter also declined from pre-surgery levels of 39 ± 1.1% to 30 ± 1.4 % at 1 – 1.5 weeks and 27 ± 1.1% at 8 weeks respectively following MI. (Fig. 3b, p < 0.0001).

**Fig. 3.**
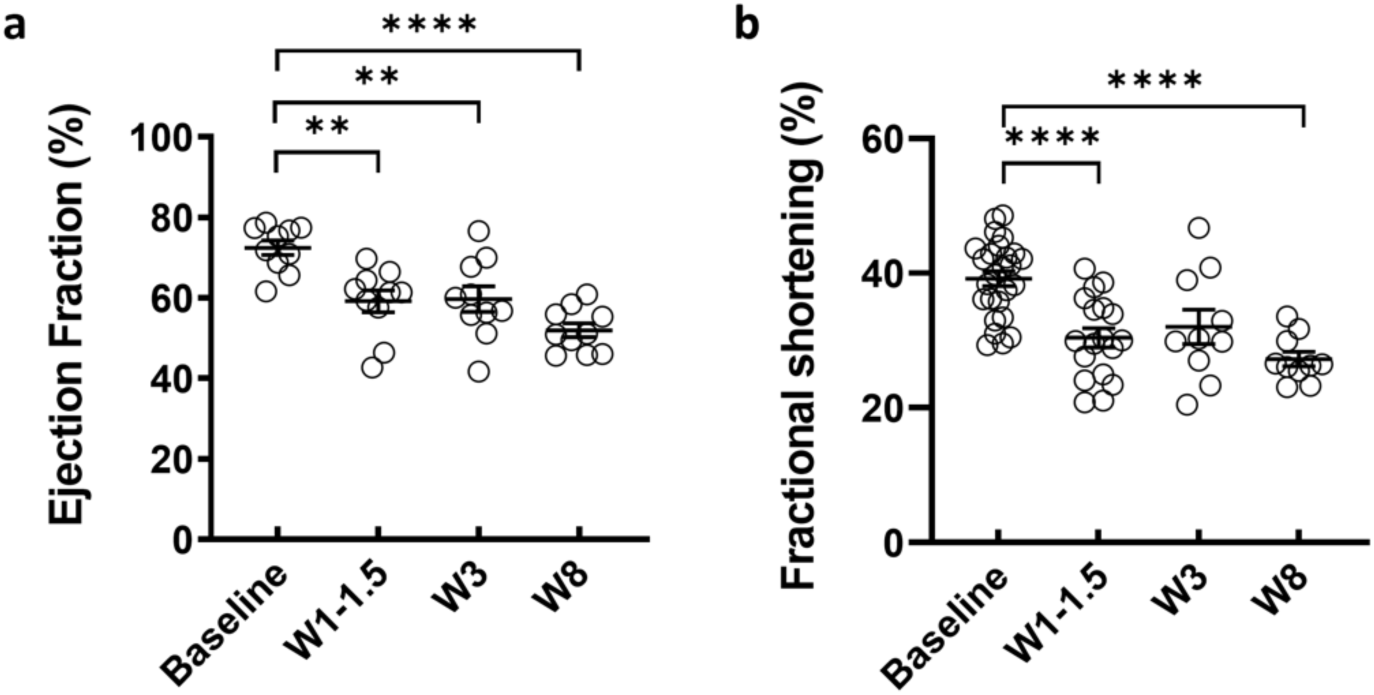
Echocardiographic evaluation of LV systolic function. Measurements of **a**, EF and **b**, FS from baseline to week 8. *N*, Baseline= 25, W1 - 1.5 = 17, W3= 10, W8 = 10. **** p < 0.0001, ** p < 0.01, compared to baseline, by mixed effects model analysis.

After an MI it is known that LV remodelling involves changes in wall thickness in both infarcted and non-infarcted regions^48–50^. We first evaluated the wall thickness of the infarcted region in comparison to the non-infarcted LV wall. Here, we present the results as the fraction of the change in wall thickness at the infarcted site when compared to the non-infarcted site on the same acquisition plane in systole and diastole. This was done to reduce the effect of variations in the acquisition plane, particularly at the distal LV level where landmarks were less clear.

At mid-LV level, there were no statistically significant changes in the wall thickness in diastole (Fig 4a) or systole (Fig 4b) following MI, which was likely due to the more apical infarct location. Conversely, at the distal-LV level, where the infarct would be more pronounced, an increase in wall thickness was noted at week 1-1.5 and week 3 in both diastole and systole compared to baseline (Fig 4c). In systole, there was an increase in wall thickness which was maintained from 1 – 1.5 to 8 weeks post MI (Fig 4d). These changes are likely due to the occurrence of eccentric hypertrophy in the non-infarcted wall, thinning of the infarcted wall segments or a combination of both. These differences were more pronounced in systole as the infarcted segment is akinetic and unable to thicken adequately compared to the hyper-contractile non-infarcted segment^51^. These wall thickness changes further support the finding of regional wall motion abnormality, which further fulfils criteria set out in the Universal Definition of MI.

**Fig. 4.**
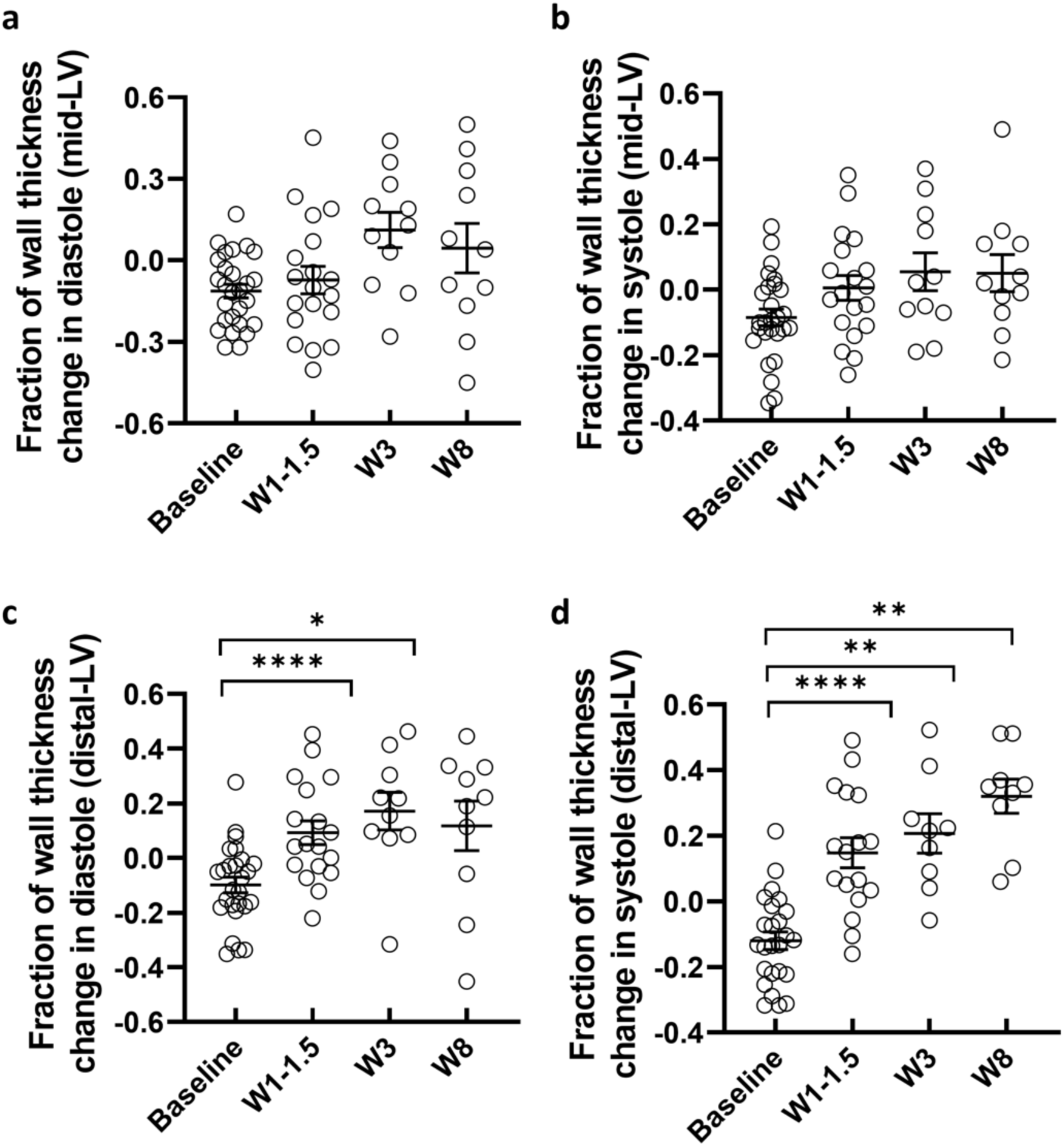
– Fraction of wall thickness change. Wall thickness changes at mid-level in (**a)**, diastole, and (**b)**, systole. *N*, baseline=27, W1-1.5=19, W3=11, W8=11. Wall thickness changes at distal-LV level in diastole (**c**) and systole (**d**) *N*, baseline=25, D3=8, W1-1.5=17, W3=9, W8=9. ****p < 0.0001, **p < 0.01, *p < 0.05 by mixed effects model analysis.

### Gross morphology and infarct size

The transient occlusion of the LAD after the second diagonal branch resulted in a well-defined area of infarction detectable at all three-time intervals despite varying appearance reflecting the acute, proliferative and maturation stages of scar formation (Fig. 5a, b and c). At 8 weeks, infarcts were more easily measured due to the presence of visible white area of scar tissue. The infarcts were less well visually defined after 3 days and 1.5 weeks. The infarct was best seen in a cross-sectional image, revealing the intra and transmural nature of scar tissue distribution. Planimetry was used to assess the size of the infarct from the anterior surface of the LV. The average size of an infarct was 4.7 ± 1 cm^2^. However, the infarct frequently extended around the apex into the posterior wall, making it difficult to accurately determine the infarct size using this measuring approach..

**Fig. 5.**
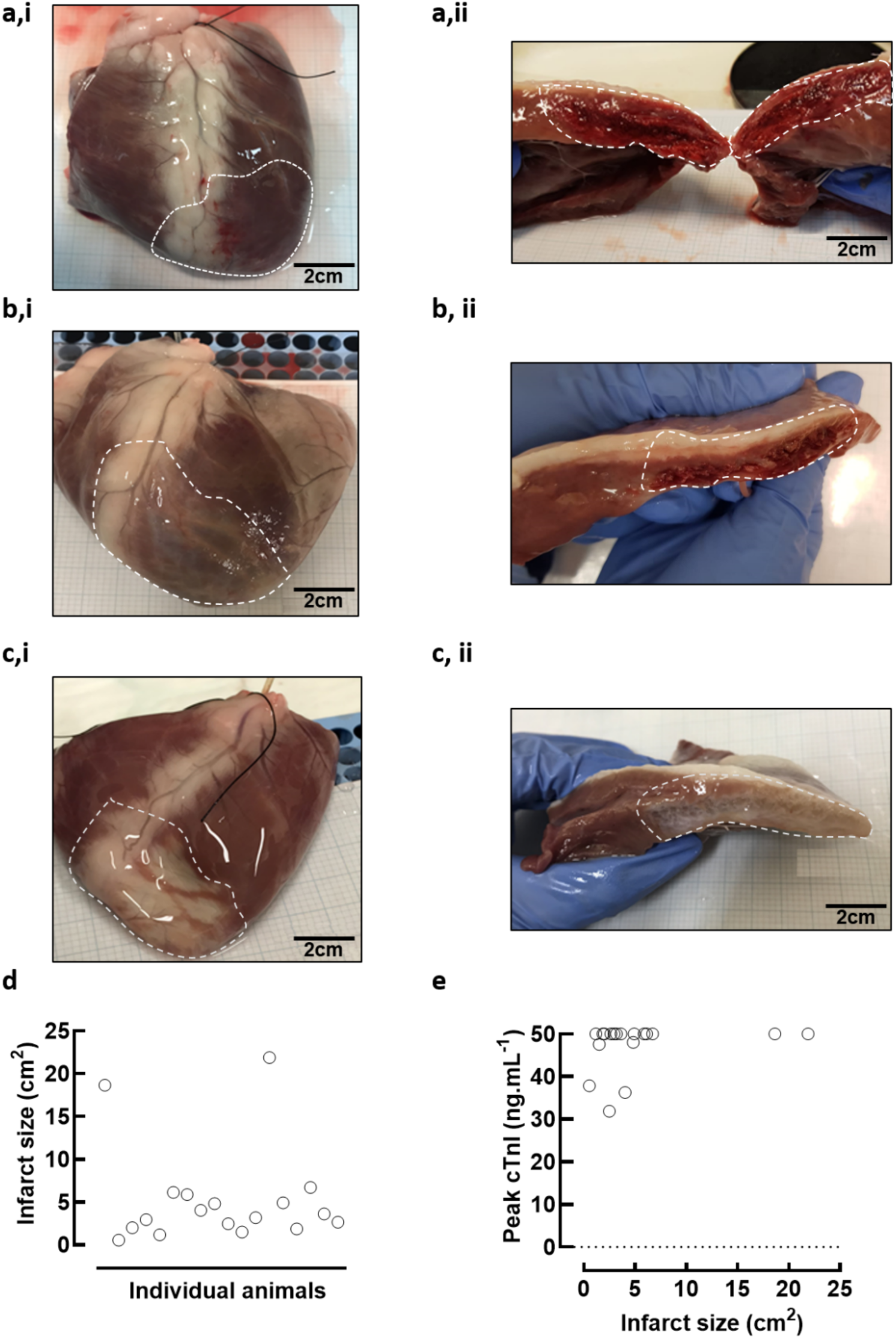
Infarcts at 3 days, 1.5 weeks and 8 weeks. Images of infarcts at **a**, 3 days, **b**, 1.5 weeks and **c**, 8 weeks as **i**, whole ventricle images and **ii**, LV cross-sectional images demonstrating different appearances in the infarct over time. The scale bar is shown in black. **d,** Infarct sizes from all animals at 3 days, 1.5 weeks and 8 weeks. **e**, The peak troponin value plotted against the infarct size with no relationship seen (*N*=26). Relationship are determined by simple linear regression.

The variability in infarct size is consistent with the variable infarct size seen in humans. No correlation between infarct size and troponin levels (Fig. 5e) were observed but this is likely related to the limitations on both infarct size measurement and the upper limits of detection of the point of care cTnI assay (50ng/ml).

## Discussion

We aimed to create a refined minimally invasive ischaemia/infarction-reperfusion model of myocardial infarction with improved survival outcomes. It was critical that our model survived to the specified time points of 3 days, 1.5 weeks, and 8 weeks in order to assess the temporal evolution of the acute, latent, and chronic phases following a myocardial infarction. In order to achieve this, the experimental protocol was refined with specific measures to reduce the mortality rate previously described in the literature^28^ including the inclusion of prophylactic anti-arrhythmic medications^47, 52, 53^ and the implantation of an internal cardiac defibrillator. Using this surgical and medication protocol, we successfully created reproducible infarcts and reduced procedural mortality. Importantly, the model fulfils the key criteria of the 4^th^ universal definition of MI demonstrating i) the rise and fall of a cardiac biomarker (cTni), ii) electrocardiographic changes in the ST-T segment, iii) echocardiographic evidence of wall motion abnormality and iv) evidence of scar/infarct on the heart.

During the occlusion period, we observed cardiac repolarization abnormalities in the form of ST-T segment changes. The presence of ST elevation indicates a transmural infarction, which affects the heterogeneity of the ionic properties of cardiac cells from the epicardial, myocardial, and endocardial layers^54, 55^. It is debateable as whether the repolarisation abnormalities are more reflective of transmural injury or the baso-apical position of the injury^56^.

We also observed changes in contractile function and wall thickness over the temporal evolution of the MI. It is known that, during the post infarct remodelling phase, the infarcted segment undergoes thinning and expansion. The apico-anterior segments are particularly vulnerable to this as they are the thinnest segments with the greatest curvature^57–60^. This is accompanied by hypertrophy of the non-infarcted region^57–62^, which may serve as a temporary compensatory mechanism. The hypertrophy occurs in response to increased wall stress as a consequence of the infarcted segment^61^. This is often not enough to restore the original function, as we have demonstrated here. In fact, this eccentric hypertrophy often contributes to worsening dilatation during remodelling^60^, which may explain to the deterioration in LV function observed at 8 weeks.

There was large inter-animal variability in infarct size with a coefficient of variation of 104%. This is likely due to the inter-animal LAD anatomical variations in the D2 bifurcation point and the limitations of the infarct measurement method. In an effort to standardize the approach, we aimed to occlude the LAD after the D2 bifurcation. However, the variability in coronary anatomy, particularly of the LAD vessel which has been previously described in sheep is likely to have contributed to this^45^. We aimed to achieve an anterior-apical infarct involving ∼ 25% of the LV. In some animals, the point of D2 bifurcation was quite proximal and would affect more than the intended area of myocardium. Whereas in other animals, the D2 branch followed a similar course to the LAD towards the apex potentially co-supplying the intended infarct territory.

Finally, we demonstrated a lower rate of attrition and improved mortality rate compared to existing work^20, 28, 29^ with the refinements in our experimental protocol. The models have survived the planned experimental time periods successfully allowing the temporal evolution of MI to be studied at an *in vivo* level and will allow future cellular and tissue studies.

Finally, we demonstrated that this model is clinically relevant due to similarities with existing STEMI patients undergoing PPCI reperfusion treatment. As a result, this model will allow for a better understanding of the pathological evolution following MI and will help in the research of new therapeutic targets that may improve patient outcomes post MI. Furthermore, with further refinement, this model may be able to reflect a translatable HF model of ischemic cardiomyopathy following STEMI and could be used to select cardioprotective medications to protect STEMI patients from reperfusion damage.

The refinement aspects developed in this study encompassed the inclusion of a prophylactic intra-operative antiarrhythmic strategy including the use of amiodarone, lidocaine and atenolol and the use of an implantable cardiac defibrillator. Whilst approximately one-third of animals developed ventricular arrhythmias requiring defibrillation, our overall intra-operative mortality was reduced to 6.7%. This compares favourably to previous studies^20, 26, 29, 37^ and provides a methodological approach that is easily applied to other large animal ischaemia reperfusion studies.

## Methods

### Ethical statement

All procedures involving the use of animals were performed in accordance with the United Kingdom (UK) Animals (Scientific Procedures) Act, 1986 and European Union Directive 2010/63. Local ethical approval was obtained from the University of Manchester Animal Welfare and Ethical Review Board. Reporting of animal experiments was in accordance with the ARRIVE guidelines 2.0 ^31^.

### Animal

Experiments were performed on naïve adult (∼18 months) female Welsh Mountain sheep. Animals were not randomised as this study focused on model development and did not contain a sham-operated arm as all statistical comparisons were paired to pre-surgical values in the same animals. Animals were group housed, fed hay and water *ad libitum*, and maintained on a 12-hour light/12-hour dark cycle for at least 1 week prior to surgical intervention.

### Myocardial infarction surgery

In line with all experiments necessitating the use of general anaesthetic, animals were fasted overnight to prevent the risks of gastric distension but had unrestricted access to water. The full step by step protocol is available in Extended Methods Online Supplement. Induction of anaesthesia was achieved using a combination of isoflurane, (5% vol/vol) Santa Cruz Biotechnology, USA), Nitric Oxide (50%vol/vol) and O_2_ (50% vol/vol at a flow rate of 5 L/min) administered via facemask. The depth of anaesthesia was confirmed by loss of the corneal blink reflex^63^. To facilitate the passage of the endotracheal tube into the trachea, lignocaine local anaesthesia spray (Xylocaine, Astra Zeneca, UK) was applied topically, reducing vocal cords and pharynx spams and tracheal intubation with a curred endotracheal tube (8-10 mm). The cuffed endotracheal tube was then inflated to ensure a sealed circuit, avoid anaesthetic leakage, and prevent secretions and gastric contents from entering the lungs. The tube was then connected to a mechanical tidal ventilator (Zoovent, UK) and ventilation performed at a rate of 15 breaths/min. The maintenance of anaesthesia was achieved using an isoflurane concentration ∼3% mixed with O_2_ (4L/min). The following parameters were constantly monitored during the surgery: i) the depth of anaesthesia was monitored by assessing the corneal blink reflex, ii) arterial blood pressure an electronic sphygmomanometer (Cardell Veterinary Monitor 9402, Sharn, USA) placed on the shaved tail base, iii) arterial O2 saturation (kept above 95%) using a doppler pulse oximeter (Nonin Medical, Inc., USA) placed on the tongue or on the shaved ear and, iv) the electrocardiogram was monitored using a five-lead continuous ECG (EMKA Technologies) digitised to a computer at a sampling rate of 1kHz (iOX2, EMKA Technologies).

In order to correct for any blood loss during the surgery, an intravenous (IV) maintenance fluid (0.9% NaCl, Baxter, USA) was administrated via an 18 to 22-gauge cannula (BD Microlance™, UK) positioned in the lateral saphenous vein from the right posterior limb at a constant flow rate over the course of surgery. This venous line was also used as a route for the administration of intravenous drugs (Fig. 6) via a coupled three-way tap.

**Fig. 6.**
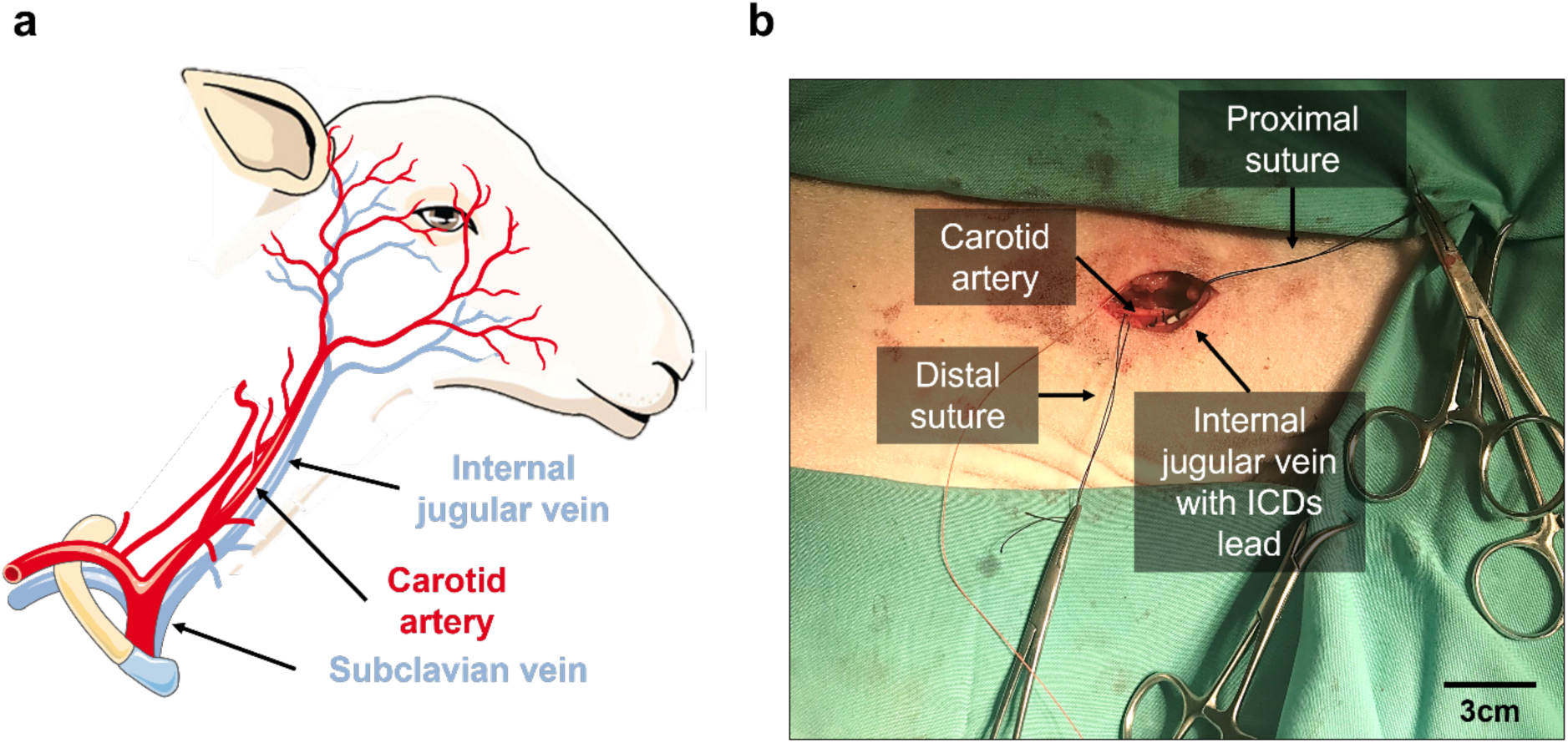
Intraoperative images of surgery. **a,** Schematic of the isolated vessels in the sheep neck. **b,** Picture of the surgical site (right side of neck) shaved and cleaned, showing the incision with sutures placed on the distal and proximal aspect of the carotid artery.

Meloxicam (0.5 mg.kg^-1^ subcutaneously; Metacam®, Boehringer Ingelheim, Germany) and amoxicillin (20µg.kg^-1^ Betamox®, intramusculalrly Norbrook, UK; or amoxicillin sodium, intramuscularly, Bowmed Ibisqus Ltd, UK) were used to provide analgesia and antibiosis. The surgery was carried out in two stages under aseptic conditions.

### Implantation of the internal cardiac defibrillator (ICD)

The first stage consists of the implantation of an internal cardioverter defibrillator (ICD) device in order to allow prompt cardioversion of any sustained intra-operative ventricular arrhythmias and record the electrical cardiac function in the post-operative period. In detail, the animal was positioned in left lateral recumbency. A sterile field was prepared by shaving the wool over the right cervical region and cleansing with iodine (iodinated povidone 7.5 % Videne, UK). An ∼ 8 cm skin incision was made in the jugular groove and bunt dissection was used out to expose the jugular vein. Once adequately freed of attached fascia, 2.0 silk sutures (Ethicon, USA) were loosely placed proximally and distally. Further blunt dissection was then carried out to identify the carotid artery, which runs deeper alongside the vagus nerve. The carotid artery was then freed from the vagus nerve using blunt dissection and 2.0 silk anchoring sutures were placed loosely at the proximal and distal points of the artery (Fig. 6 a and b).

After tying off the proximal part of the jugular vein, a small incision was made into the vein and maintained open using a vein pick to allow the insertion of an ICD lead (Medtronic USA) The single chamber defibrillator lead comprises, from proximal to distal, an electrode to be anchored within the myocardium, and bifurcated (in a single coil lead) or trifurcated (in dual-coil lead) header connector pins. The connectors consist of one pace-sense IS-1 (or IS-4) connector, and one or two DF-1 (or DF-4) high voltage connectors^64^. Consequently, the active fix right ventricular (RV) lead was carefully advanced to the apex of the RV under fluoroscopic guidance (BV Pulsera, Philips, Netherlands).

The correct lead tip position was assessed prior to fixation in the RV by connecting the lead to an electrophysiological analyzer (Medtronic 209o Analyser; Medtronic Inc) and assessing three intraoperative electrocardiographic parameters: (i) the pacing threshold (i.e., the lowest pulse amplitude at which the heart can still be paced); (2) intracardiac potential (R-wave amplitude), and (3) lead impedance (300 - 1500Ω)^65, 66^. Satisfactory elad positioning was confirmed by a combination of i) pacing threshold < 1V, ii) R wave amplitude > 6mV and lead impedance < 1500 Ω. Once satisfactory lead and pacing parameters were confirmed the ICD lead was connected to the ICD can and secured in position using 2.0 silk suture where it entered the jugular vein. The latter minimizes the risk of dislodgment. Any excess lead length was wrapped in loose loops around the ICD can before being positioned inside a subcutaneous pocket created above the jugular distally to the original incision towards the right shoulder. The pocket was securely closed using Vicryl 2.0 sutures. To avoid inappropriate shocks, the device was configured to only detect the following zones: VT, VF and Fast VT zones). In this ovine model, inappropriate detection has been observed as high sinus tachycardia rates and T wave over-sensing, therefore the device was programmed purely for detection. The defibrillator is then set to emergency mode for the duration of the surgery, allowing for fast defibrillation when intra-operative ventricular arrhythmias.

### Induction of myocardial infarction

Access to the arterial circulation was obtained via the carotid artery which was identified at the start of the surgery. The originally placed 2.0 suture at the proximal end of the vessel was secured. Using the Seldinger technique^67^, the carotid was punctured with a 14-gauge cannula. A soft curved tip guidewire (Abbott, USA) was then inserted through the cannula and advanced into the arterial lumen. The guidewire was securely held in place whilst the cannula was withdrawn. A 12 cm 6Fr haemostatic introducer sheath (Abbott, USA) was passed over the guidewire into the lumen. The guidewire was withdrawn, leaving the introducer sheath in the vessel. The sheath was loosely secured with a suture placed into the surrounding soft tissue to avoid displacement due to carotid pulsation. 10,000 IU of heparin sodium (Wockhardt Ltd., India) was injected as a bolus, and the IV maintenance fluid at the right posterior limb cannula was switched for one containing 0.9% NaCl IV infusion at 10 IU heparin per mL. The medication protocol including the prophylactic anti-arrhythmic drug regime is demonstrated in Fig. 7.

**Fig. 7.**
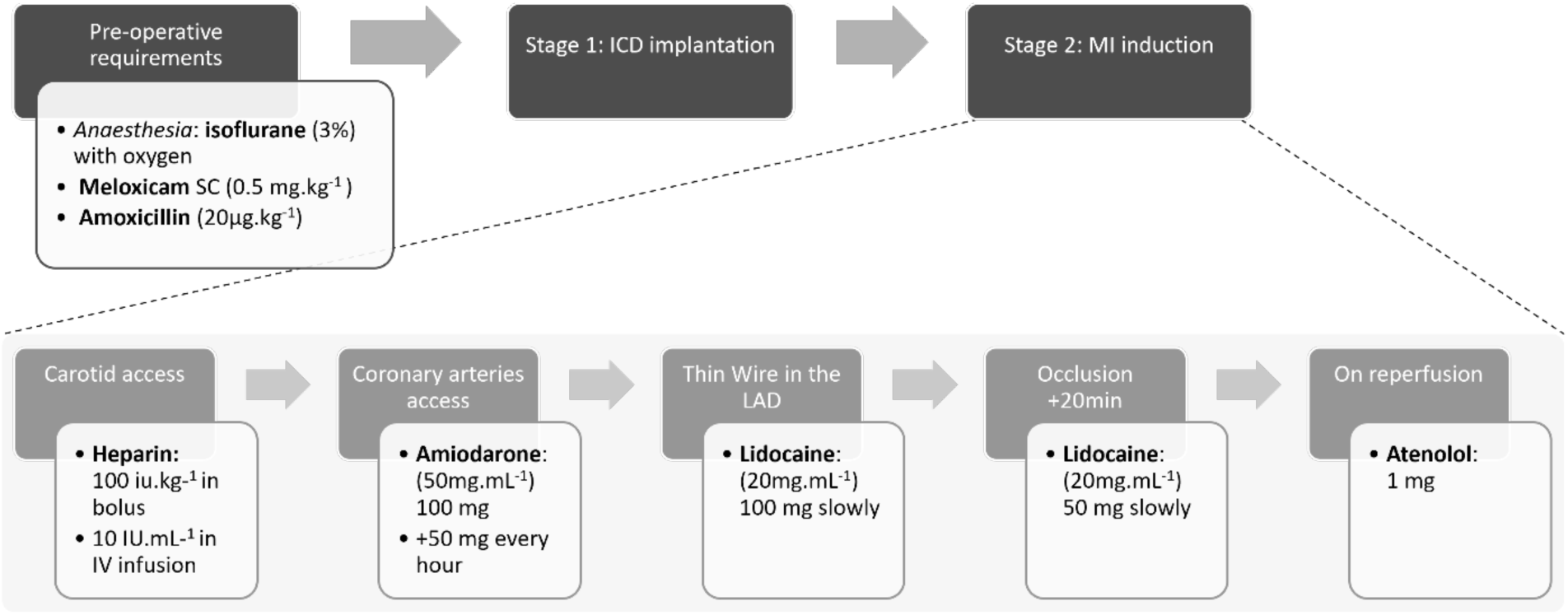
Drug protocol for MI induction surgery. Diagram of the timings of medications administered during MI induction surgery. Internal cardiac defibrillator (ICD) device; anterior descending artery (LAD).

A 6Fr Judkins right (JR4) catheter (Runway, Boston Scientific Quincy, USA) was preloaded with a 0.35mm J tip wire (Cordis, USA or Abbott, USA) and advanced through the introducer sheath to the aortic valve level under fluoroscopic guidance leading with the J-tipped wire.

The wire was then removed, and the left main stem (LMS) was engaged with a 50:50 mix of radiocontrast agent (Visipaque^TM^, GE Healthcare Inc., USA; 270 mg/mL) and 0.9% NaCl solution injected through the sheath catheter. A coronary angiogram was performed to opacify the left coronary system. The left coronary system consisted of the left main stem which bifurcates into the left circumflex (LCx) and left anterior descending (LAD) coronary artery, also known as the left homonymous in sheep ^17, 61–63^. The LCx supplies the posterior and lateral free walls of the left ventricle (LV) whereas the LAD supplies the anterolateral, septal and apical walls of the LV^62, 64, 65^. Upon identification of the LAD, a hydrophilic 0.014-inch guidewire (Abbott, USA) was advanced to the distal LAD. Depending on the calibre of the vessel, an appropriately sized intracoronary balloon catheter (Apex Monorail or Emerge^TM^, Boston Scientific, MA, USA) was advanced and positioned within the LAD, after the second diagonal branch (D2; Supplemental Figure SI). The length of the intracoronary balloon used was 20 – 40 mm with a width ranging from 2 - 2.75 mm depending on the coronary anatomy and calibre determined empirically during coronary angiography using the Judkins catheter diameter as a guide. The balloon catheter was inflated to the specified recommendations using an Indeflator (BasixCompak inflation device, Merit Medical, USA) with a repeat coronary angiogram thereafter confirming complete occlusion of flow distal to the inflated balloon (Fig. 6; Supplemental Figure SI). The balloon remained inflated for 90 minutes occluding blood flow distal to the balloon, thus creating the infarct.

ECG and intracardiac electrogram (via ICD) parameters were continuously monitored for the presence of ST elevation, T wave changes and conduction abnormalities and ventricular arrhytmias. ST segment changes were usually observed within minutes of coronary occlusion with evidence of additional ventricular activity seen approximately 30 - 40 minutes into the occlusion. These ranged from the occasional ventricular ectopic (VE) beats to VF requiring defibrillation. In the event of a persistent ventricular arrhythmias such as VT or VF, the animal would be promptly defibrillated via the ICD to terminate the arrhythmia.

90 minutes after inflation the balloon was deflated and coronary blood flow distal to the occlusion site was confirmed with a repeat coronary angiogram Fig. 8. The intracardiac equipment and sheath were removed. The distal carotid artery was tied off with a 2.0 silk suture. The wound was closed in three layers, closing the muscular fascia, the subdermal tissue and finally the cutaneous layer. During the wound closure, the isoflurane concentration was gradually reduced from 3% to 0%. The endotracheal tube was left in place until the animal showed the signs of a rejection reflex (i.e., swallow or cough). The recovery from anaesthesia was closely monitored for evidence of respiratory distress and arrhythmias. The IV catheter was left in place until the animal was fully awake to allow immediate IV access. Once the animal was alert and all vital signs were within normal range, it was placed in a single housed post-operative recovery pen in full sight and communication with its peers and was given access to food and water.

**Fig. 8.**
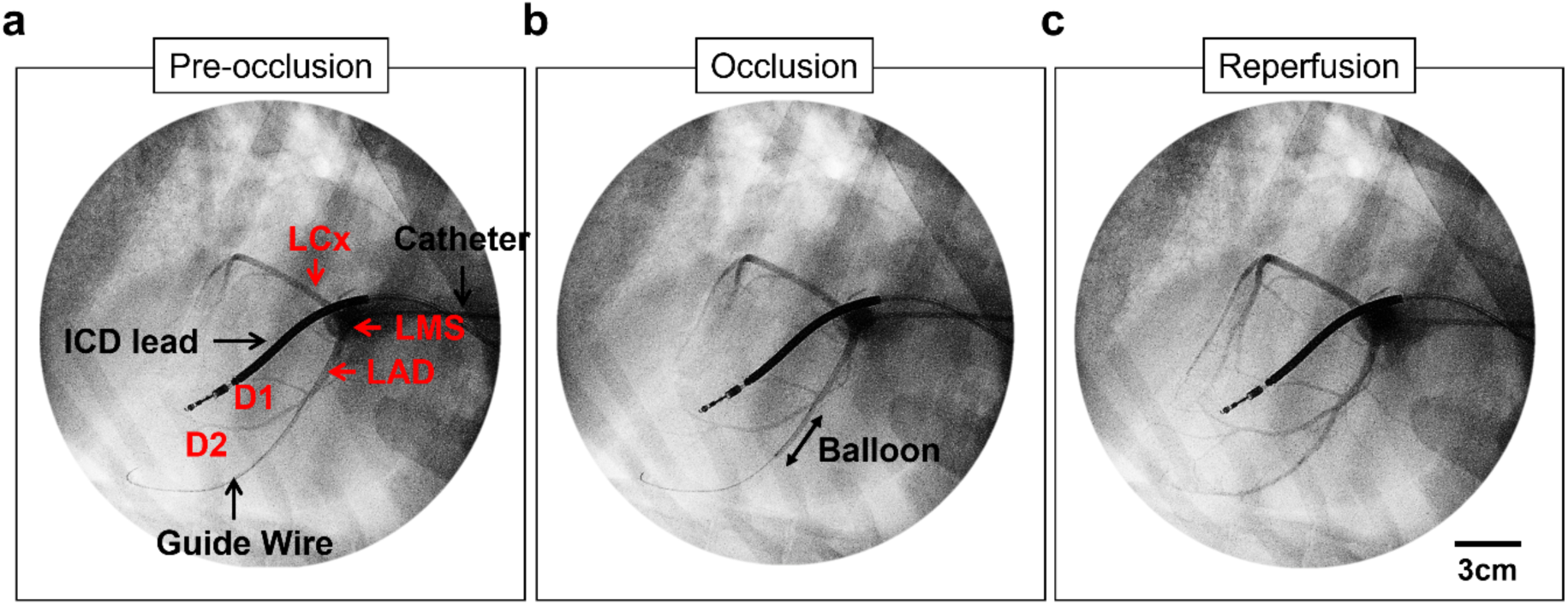
Fluoroscopic images during MI induction surgery. **a**, Angiogram of the left coronary system showing the left main stem (LMS) bifurcating into the left anterior descending (LAD) and left circumflex (LCx) coronary artery. The ICD lead is also seen within the RV. **b,** Repeat coronary angiography with an inflated intracoronary balloon occluding flow distal to the balloon. **c,** Confirmation of the correct reperfusion of the heart. D1 and D2, diagonal branches. RA, right atria. RCA, right common coronary artery. Videos in supplemental material (Supplemental material SI)

The animal was closely monitored during the recovery period. It was considered fully recovered from anaesthesia when visibly alert, standing and having urinated. After 24h of post-operative recovery housing the animal was returned to group housing. No further intervention was carried out for at least three days to ensure complete recovery from surgery as well as to allow the ICD lead to settle in its intracardiac position. Animal weight, well-being and wound healing were monitored regularly to ensure adequate post-operative recovery.

### *In vivo* assessments

To assess the evolution of various clinical and cellular parameters over time following MI, the animals were randomly divided into three groups: 3-day MI, 1 - 1.5 week MI, and 8 week MI. As shown in Fig. 9, *in vivo* measurements were performed at baseline, post-operatively, and at the endpoint. There were two types of evaluations performed: full assessments and expedited assessments. The full in vivo assessment included measurement of weight and blood pressure, recording of electrocardiograms (ECG), imaging using transthoracic echocardiography (TTE), blood sampling, and external interrogation of an intracardiac device once implanted. The expedited assessment only involved blood sampling and the device interrogation.

**Fig. 9.**
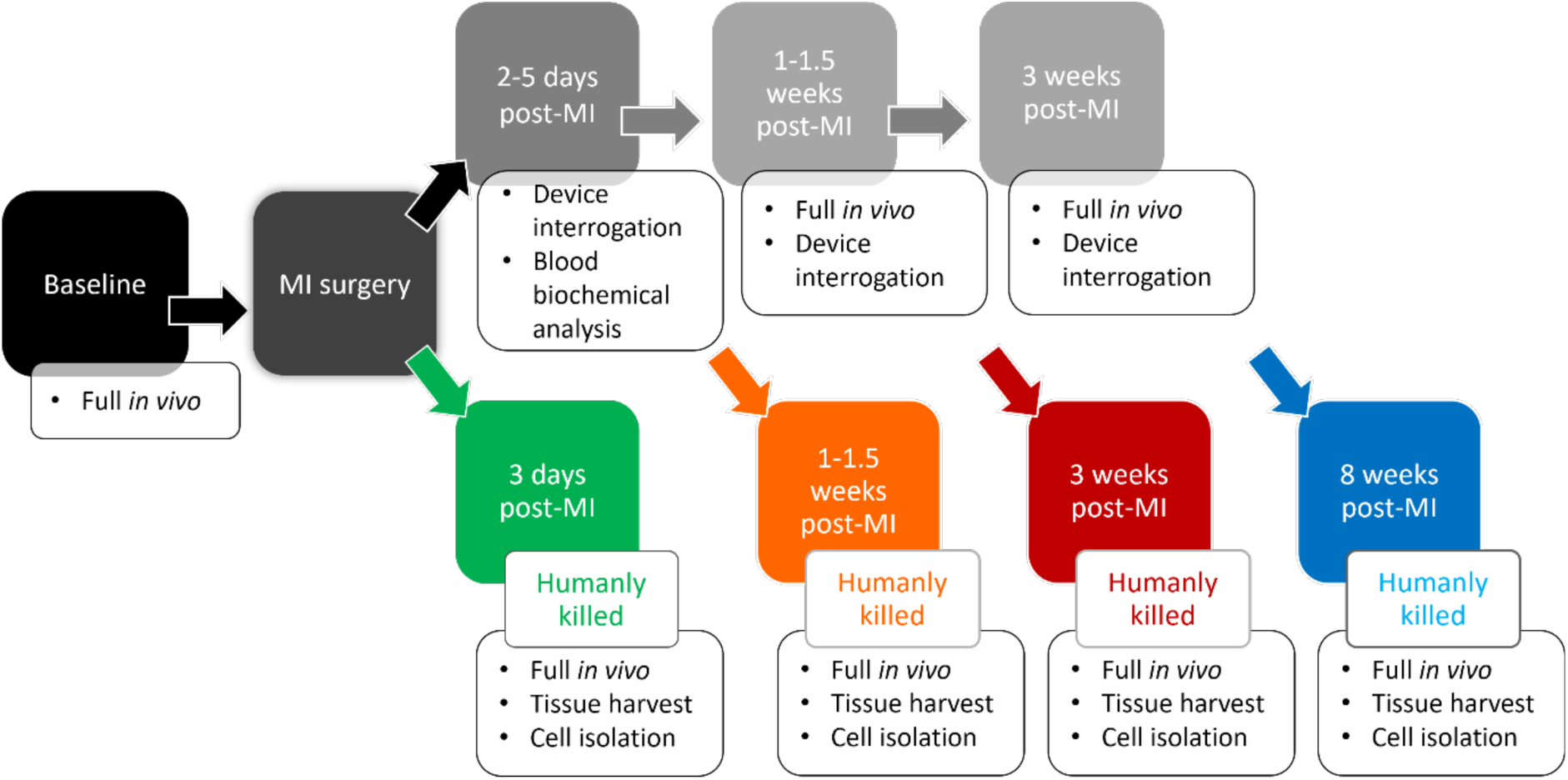
*In vivo* schedule for 8 week, 3 week, 1-1.5 week and 3 day MI animals. The full *in vivo* assessment included measurement of weight and blood pressure, recording of electrocardiograms, imaging using transthoracic echocardiography, blood sampling, and external interrogation of an intracardiac device once implanted.

### Blood sampling

Blood was collected pre-operatively, at multiple time points intraoperatively (carotid artery access, pre-occlusion, 30min post-reperfusion and 90min post-reperfusion), at 2 - 5 days, 1 - 1.5 weeks and 8 weeks. Venous blood sampling was performed preferably from the right jugular vein with the animal gently restrained using a sterile aseptic non-touch technique. Alternative sampling sites included the left jugular vein or cephalic veins. The skin was shaved and aseptically cleansed before sampling. Amongst the blood parameters evaluated were whole blood cTnI (VetScan i-STAT, Abaxis, UK) as a marker of myocardial necrosis along with a full biochemistry profile (Skyla VB1 Analyser, Woodley, UK). The full profile is shown in Supplement (Supplemental Table 1). The cTnI was tested using a point-of-care testing system, which provided a result within 10 minutes.

### Transthoracic echocardiography

In the same seated position, under gentle restraint, transthoracic echocardiography (Vivid, GE Healthcare,USA) was performed with the probe positioned on the right side of the chest with the right forelimb lifted, providing access to the thorax. Image quality was optimised by shaving the chest and using an ultrasound transmission gel (Aquasonic, Germany) for better contact. As limited views were available, parasternal short (SA) and long (LA) axis views were obtained to allow evaluation of the cardiac structure and function. For adequate visualization of the LV walls and endocardial borders, the apical views (2, 4 and 5 chamber views) would have been ideal. However, the anatomical position of the heart and wide sternum in sheep makes this difficult to obtain^56^. The SA-mid view was taken below the mitral valve level where the papillary muscles were visible. The SA-distal view was the most distally obtainable short axis image of the LV to visualize the apex and adjacent region. In humans, regional wall motion abnormalities can be assessed using a 17-segment ECHO model where the LV walls are divided into 17 segments which can be correlated to the blood supply^57, 58^.

For the analysis, images were exported and converted from DICOM to JPEG format using Microdicom Viewer (Microdicom, Bulgaria). The JPEG files were opened in Fiji ImageJ (National Institutes of Health, USA) and multiple measurements were taken.

In the SA-mid and SA-distal images, the anterior wall thickness measurement represented the infarcted region, and the posterior wall thickness represented the non-infarcted region (Fig. 10 and Fig. 11). Prior to the surgery, measurements were also taken from these identical sites to serve as baseline values. The fraction of wall thickness change (FWTC) was calculated using Equation 1 to determine the difference in wall thickness between the infarcted and the non-infarcted region. In order to assess the fractional area change (FAC) as a measure of contractile function, the LV cavity area (Fig. 10 and Fig. 11) was also measured at mid and distal LV levels in systole and diastole and calculated using Equation 2.Equation 1

**Fig. 10.**
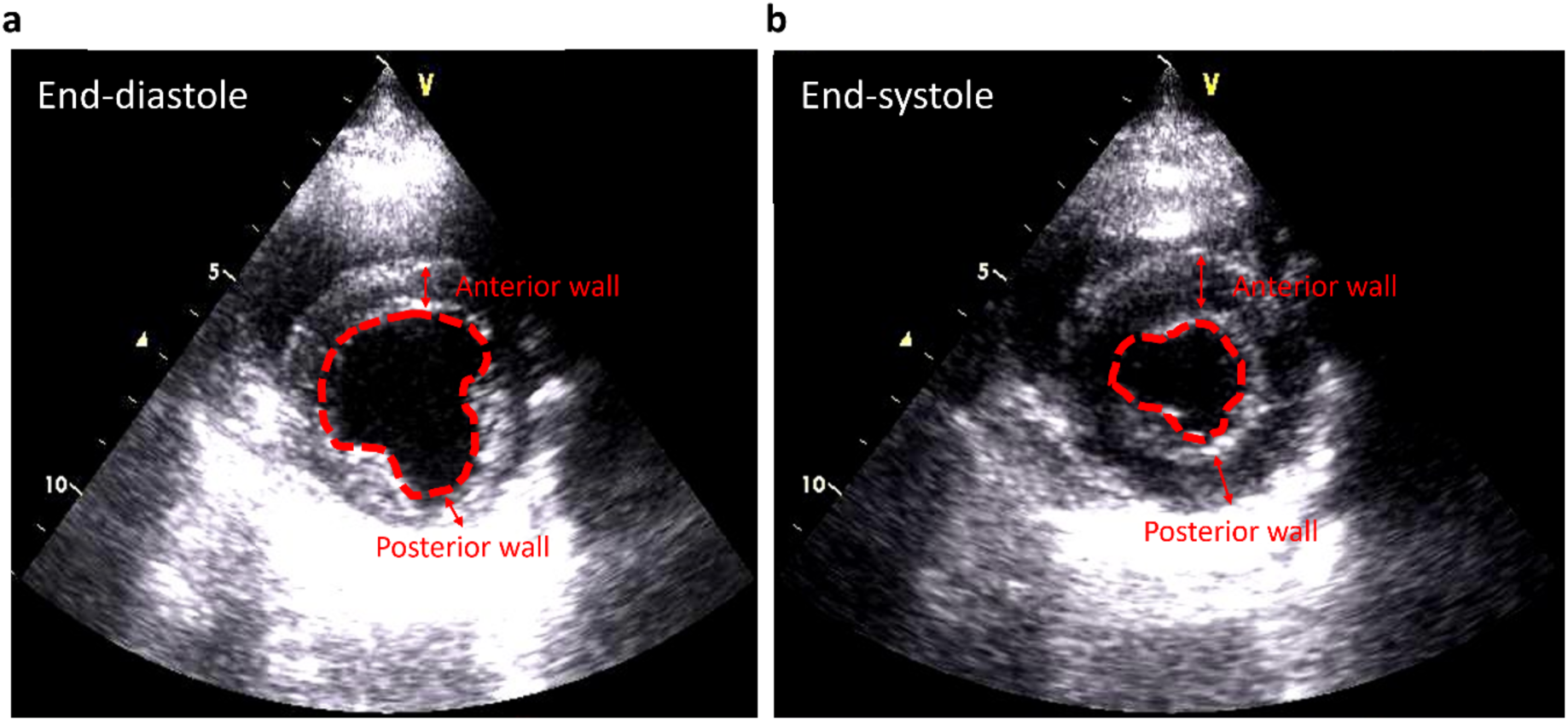
Measurements from ECHO SA-mid views. Frames from the LV short axis views at mid-level in diastole **a,** and systole **b,** with the anterior and posterior wall thickness measurement sites (marked by red arrow) as well as the LV cavity area measurement (outlined in red-dotted line) shown.

**Fig. 11.**
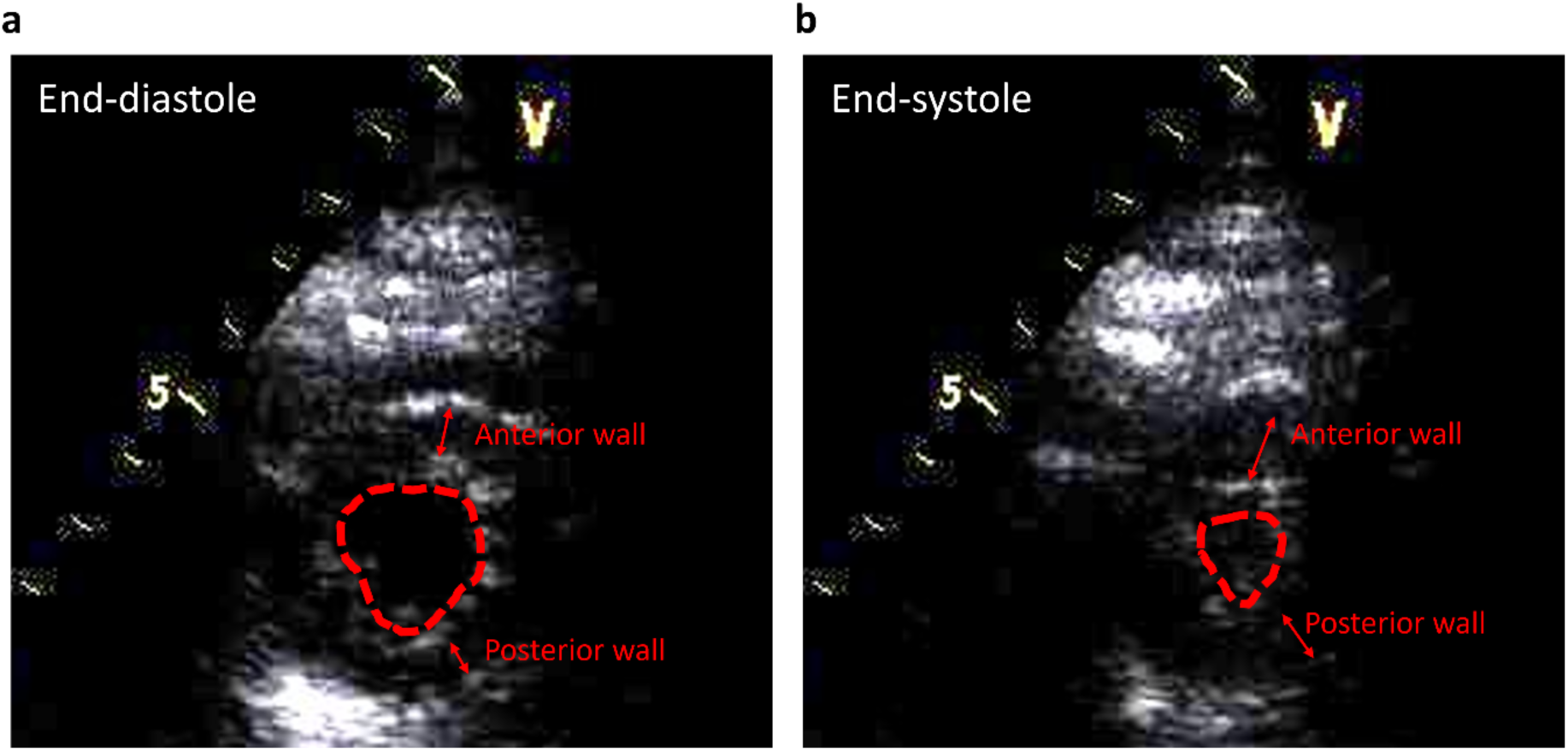
Measurements from ECHO SA-distal views. Frames from the LV short axis views at distal-level in diastole **a,** and systole **b,** with the anterior and posterior wall thickness measurement sites (marked by red arrows) as well as the LV cavity area measurement (outlined in red-dotted line) shown.

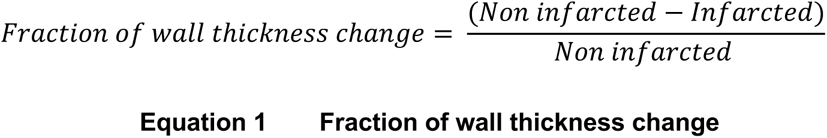

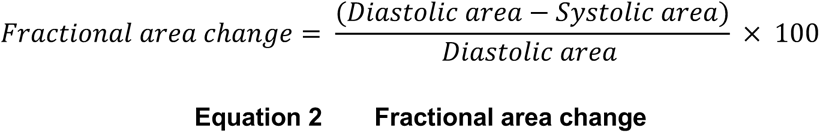

From the parasternal long axis (PLAx) images, the interventricular septal (IVS) thickness, LV end diastolic (LVEDD) and systolic diameter (LVESD) as well as the LV posterior wall (LPW) thickness were measured in diastole and systole as shown in Fig. 12. Due to the limited obtainable views in the sheep, estimating the LV function was challenging particularly in the presence of an apical infarct. Therefore, an adaptation of the simplified Quinones equation produce by MD Math available from the Canadian Society of Echocardiography^59^ was used deriving an estimated ejection fraction (EF) from the LVESD, LVEDD and an apical contractility estimate seen from the SA-distal views using Equation 3. In this equation, the K value (Table represented the apical contractility. Additionally, fractional shortening (FS) was also calculated as another measure of contractile function using Equation 4 and expressed as a percentage.

**Table 1.** K values. Apical contractility measurements expressed as percentages, used in Equation 2.4

| Apical contraction | K value (%) |
| --- | --- |
| Normal | +10 |
| Hypokinetic | +5 |
| Akinetic | 0 |
| Dyskinetic | -5 |
| Apical aneurysm | -10 |

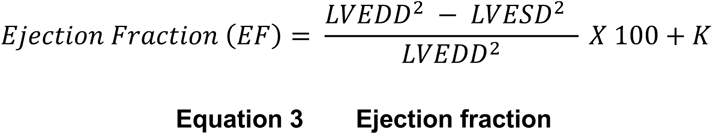

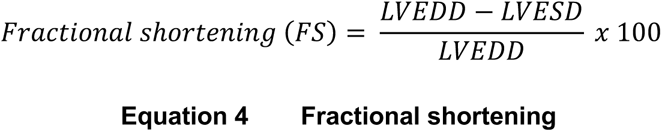

**Fig. 12.**
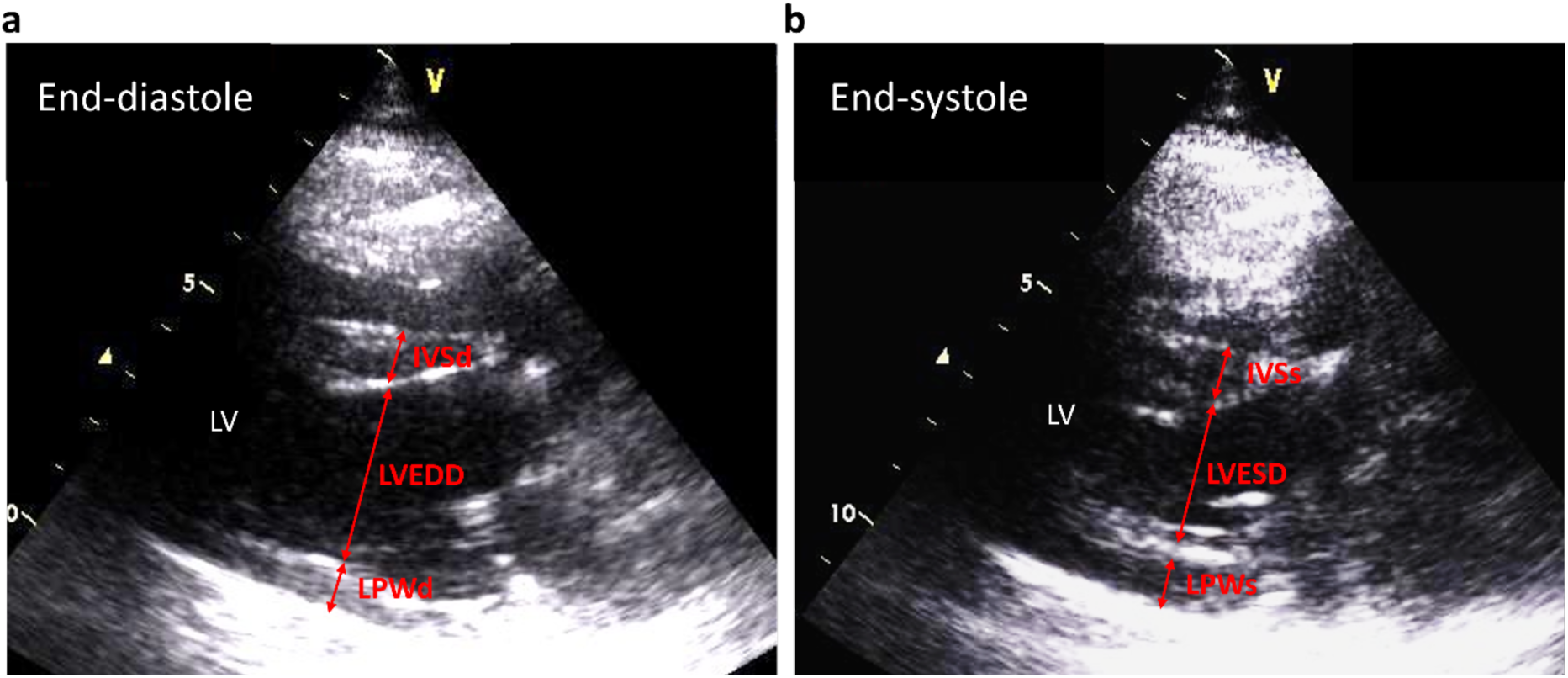
Measurements from ECHO PLAX views. Frames from the PLAX views in diastole **a,** and systole **b,** with the interventricular septal thickness (IVS) (yellow arrows), LV end diastolic (LVEDD) and systolic diameter (LVESD) (blue arrows) and left posterior wall thickness (LPW) (orange arrows). For IVS and LPW measurements, the letter *s* or *d* at the end denote measurements taken in systole or diastole, respectively.

### Electrocardiography

Surface electrocardiograms were recorded at intervals. Peri-operative ECG recordings (EMKA Technologies, USA) were performed in the seated position using 5 surface electrodes in an orthogonal arrangement on shaved and cleaned sites to allow better contact. The identical electrode position is used again intraoperatively but the animal is laid on its left-hand side during surgery. Upon connecting electrodes, the animal was allowed some time to settle thus minimizing any potential stress-related ECG changes^60^ or motion artefacts. A 10-minute ECG was then recorded through IOX software (EMKA Technologies, France)

For the analysis, the files were opened in ECG Auto (EMKA Technologies, France) and converted text file (*.TXT) format to allow the files to be opened on Labchart (AD Instruments, UK). The recordings were smoothed and filtered to reduce noise and breathing artefacts using a 21-point (Bartlett) window and a 2 Hz high pass or 50 Hz notch filter.

The software generated multiple averaged traces consisting of 10 consecutive beats over the 1-minute selected recording period where intervals could be manually selected.

### Euthanasia

The humane killing of MI sheep was carried out as per the approved regulated procedure and schedule 1 protocol in accordance with ABPA regulations with an anaesthetic overdose (pentobardbitone sodium, 200 mg/kg intravenously) followed by permanent cessation of circulation. Humane killing was performed at three predetermined timepoints post MI: 3 days, 1.5 weeks and 8 weeks post-MI.

### Statistical analysis

All data is expressed as the mean ± SEM. Statistical analysis was performed using Prism 7 (GraphPad Software, San Diego, California). Data were first tested for normality using Shapiro-Wilk, Kolmogorov-Smirnov, Anderson-Darling, and D’Agostino-Pearson omnibus in GraphPad Prism. Where data were not normally distributed, data were transformed using, natural log, Log10, reciprocal, square root or exponential depending on skew^67^. Normally distributed data were analysed using the t-test, one way ANOVA, repeated measures one way ANOVA and mixed effects model analysis. Where data was not normally distributed despite transformation, an equivalent non-parametric test was used. The relationship between two variables was determined using simple linear regression and the correlation was evaluated using Pearson’s correlation (provided by the r value). Data was considered significant if the p value was < 0.05 and is described in the text.

### Data Availability

The data supporting this manuscript are available from the corresponding author on reasonable request.

## Acknowledgements

The authors wish to acknowledge support from The British Heart Foundation (IG/15/2/31514; FS/17/52/33113 & FS/20/6/34990) and Medical Research Council (MR/K50028231/1).

For the purpose of open access, the author has applied a Creative Commons Attribution (CC BY) licence to any Author Accepted Manuscript version arising from this submission.

## Ethics Statements

The authors have no financial or competing interests to declare

## Author Information

AWT conceived the experiments and secured all funding. CP led the model development and performed all analyses. All authors contributed to surgical and in vivo procedures. CP, BN and AWT drafted the manuscript. All authors approved manuscript contents.

## Step by step guide

## MATERIALS

### Animals

• Young, treatment naïve Welsh Mountain sheep (aged ∼18 ± 6 months) with an average weight 38.5 ± 1.2kg
• Group housed, fed hay and water ad libitum and maintained in a 12 hour light/12 hour dark cycle for a minimum of 1 week prior to surgical intervention.

### Reagents

• Lignocaine local anaesthesia throat spray (Xylocaine, Astra Zeneca, UK)
• x3 Heparin 10,000 IU in 10 ml vials (Wockhardt, UK)
• Contrast medium – Ionohexol (Omnipaque 300, GE Healthcare, USA)
• x3 Sodium chloride 0.9% 500mls and 1000mls (Baxter, USA)
• Prophylactic antibiotics (Amoxicillin 15 mg kg^-1^) (Norbrook, UK)
• Prophylactic analgesia meloxicam (0.05mg kg^-1^) (Norbrook, UK)
• Amiodarone (Hameln, UK)
• Lidocaine (Hameln, UK)

### Premedication and Anaesthesia

• 1-chloro-2,2,2-trifluoroethyl difluoromethyl ether (isoflurane, Santa Cruz Biotechnology, USA)
• Oxygen and nitrous oxide (50:50) mix (BOC Healthcare, The Linde Group, Germany)
• Cone face mask rubber large

### Equipment

• Hypodermic needle variable sizes 18, 20, 22G (BD Microlance, UK)
• 5ml, 10ml, 20ml and 50ml syringe (BD Plastipak, UK)
• IV infusion set (CareFusion, USA)
• 14G & 20G cannula (BD Venflon, USA)
• Blood pressure cuff and machine (Mindray, Australia)
• Pulse oximeter sensor (Mindray iMEC8 Vet Manuals, Mindray Bio-Medical Electronics Co., China)
• 5 lead ECG cable with crocodile clips and recording equipment, IOX software (EMKA Technologies, France)
• iSTAT Vetscan Hand held analyser and cTni cartridges (Abaxis, UK)
• Ultrasound transmission gel (Aquasonic, Germany)
• GE Vivid 7 echocardiography machine with 5S cardiac transducer (General Electrics, USA)
• Skyla VB1 Biochemistry analyser (Woodley, UK)
• C-arm fluoroscopy machine (BV Pulsera Mobile C-arm, Philips, UK)

**Intubation** (see Fig. 13 – Typical surgical equipment. a, Intubation equipment. b, Surgical tray.

• Miller laryngoscope blade size 4
• Stiff endotracheal stylet
• Cuffed endotracheal tube (size 8.5 to 10; J.A.K Marketing, UK)
• 20mL syringe (BD Plastipak, UK)
• The ribbon to tie the ET tube in place
• Bag valve mask

### Monitoring and anaesthesia

• Anaesthetic machine (Zoovent, UK)
• Five-lead electrode cable with leg strips
• Electrocadiogram monitoring (IOX software, EMKA technologies,USA)
• Pulse oximeter sensor (Mindray iMEC8 Vet Manuals, Mindray Bio-Medical Electronics Co., China)
• Blood pressure cuff sized as tail cuff and BP recording equipment (Mindray, Australia)

### Surgical site preparation, generic operative equipment & operator preparation

• Sheep clippers
• Iodinated povidone 7.5% (Videne, UK)
• Sterile drapes

Surgical tray (see (see Fig. 13 – Typical surgical equipment. a, Intubation equipment. b, Surgical tray.)

• Gallipot
• 2.0 Vicryl sutures (Ethicon,USA)
• 2.0 Silk sutures (Ethicon,USA)
• 2.0 Monocryl sutures (Ethicon,USA)
• Sterile gauzes
• Internal cardiac defibrillator

ο Generator (Medtronic, USA)
ο Right ventricular active fixation defibrillator leads (DF1 or DF4) (Boston Scientific, Medtronic, USA)
ο ICD compatible programmer with analyser cable and header (*e.g*. Medtronic 2090, Medtronic, Minnesota, USA)
ο PSA cables
• MI Induction

ο 14G cannula (BD Venflon,USA)
ο 12cm 6Fr haemostatic introducer sheath (Abbott Medical, UK), containing the sheath, a dilator and a mini-guidewire
ο 6F JR4 Guide catheter (Runway, Boston Scientific, USA)
ο Haemostasis valve (Honor,Merit Medical, USA)
ο Indeflator (balloon inflation device) with pressure monitor (BasixCompak inflation device, Merit Medical, USA)
ο 50ml syringe (BD Plastipak, USA)
ο Intracoronary balloon catheter, variable sizes 2.2 to 2.75mm diameter, 20-40mm length (Apex Monorail, Boston Scientific, MA, USA)
ο 0.35 J tipped guide wire (Cordis, USA)
ο 0.0014 intracoronary guide wire (Abbott, USA)

**Fig. 13.**
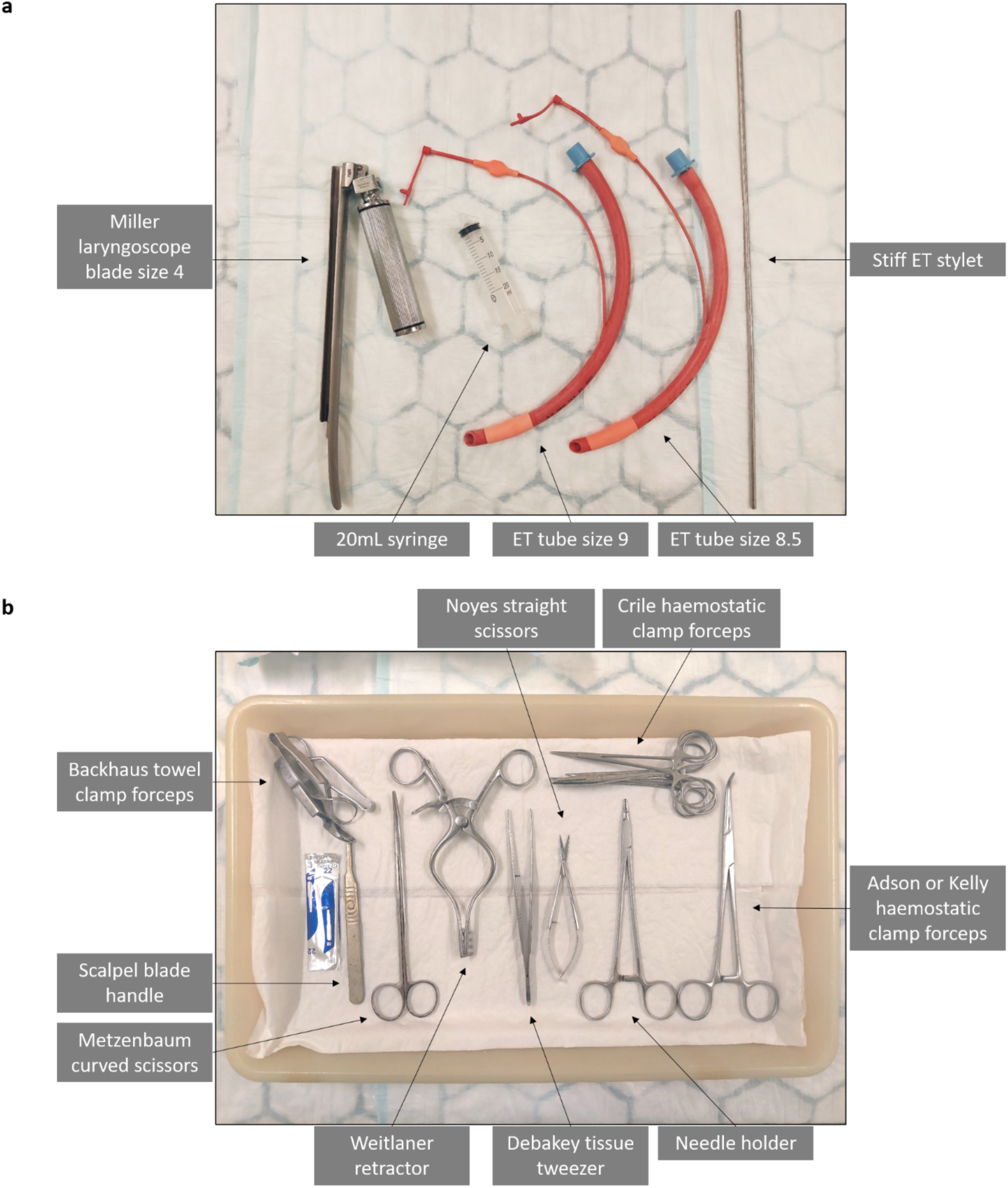
Typical surgical equipment. **a**, Intubation equipment. **b,** Surgical tray.

### Operator preparation

• Surgical gown
• Surgical cap
• Sterile gloves
• Surgical face mask
• Radiation protective equipment – lead gown (Kenex, UK) and thyroid collar

### Software

• ECG recording software details

### Surgical procedure

#### A. Anaesthetic induction and preparation

△ Prior to surgery, animals are given access to water but food is witheld to avoid ruminal distension.

1. The animal is allowed to inhale a combination of oxygen and nitrous oxide in a 50:50 mix with isoflurane (3-5%).
2. Animal is lifted and placed on the operating table in a sternal recumbency position, with the head elevated.
3. With the aid of a second operator, the jaw is opened with the head supported to allow the primary operator to spray two puffs of local anaesthetic (Lidocaine throat spray) into the back of the throat.
4. The facemask delivering oxygen nitrous 50:50 with isoflurane is replaced in preparation for intubation.
5. The facemask is removed and the jaw is held open with the head supported by the second operator. A laryngoscope is introduced and the vocal cords visualised.
6. A stiff stylet is taken down to the level of the vocal cords. An appropriately sized endotracheal tube is advanced over the stylet past the vocal cord and the stylet is removed. The cuff is inflated with air using a 20mL syringe and the tube is connected to a bag mask to confirm the appropriate tube placement in the trachea. The tube is then connected to the ventilator. The appropriate tube placement is confirmed by appropriate chest wall movement with the ventilator, saturation recordings, and the maintenance of anesthesia, and the tube is secured with the tie around the jaw.

△ When advancing stylet, due care is taken not to advance the stylet too far to avoid damage to soft tissue or larynx. Adequate visualisation of vocal cords is required to ensure safe placement.

7. The animal was positioned in lateral recumbency for the rest of the procedure.
8. All monitoring equipment (saturations monitoring, ECG monitoring, and BP monitoring) is connected.
9. A 20G cannula is sited on the right hind leg for the purposes of administration of intraoperative medications. Pre-operative antibiotics and analgesia are administered at this stage.
10. The right side of the neck is shaved, providing a wide surgical field.
11. The skin is cleaned twice with an iodine-based antiseptic and draped.
12. Anatomical landmarks are palpated to delineate the jugular groove.
13. A 5 to 7cm incision is made to the skin with a blade within the jugular grove. The incision site is located two thirds of the way from the angle of the jaw to the shoulder tip. This is followed by blunt dissection down to identify the jugular vein.
14. The jugular vein is identified and freed. A proximal and distal 2.0 silk suture is placed loosely on the vessel. These ties are clipped onto the drapes using the Crile (?artery) forceps.

△ Gentle dissection is performed to free the jugular vein and carotid artery to avoid vascular damage. The vagal nerve must be gently freed from the carotid artery.

15. Further blunt dissection is performed deeper to identify the carotid artery, which runs alongside the vagus nerve. The carotid artery is freed from the vagus nerved and similar proximal and distal 2.0 silk sutures are placed and clipped to the drape loosely.

△ The procedure is carried out in two phases. The first stage involves implantation of an internal cardiac defibrillator to manage intraoperative life-threatening ventricular arrhythmias and the second stage is the induction of MI.

#### B. Implantation of an internal cardiac defibrillator (ICD)

1. The proximal jugular vein suture is tied off.
2. The mobile C-arm of the fluoroscopy machine is moved into position over the heart in the postero-anterior position.
3. With Noyes scissors, a venotomy is performed, exposing the inner lumen of the vessel.
4. With the aid of a vein pick, the vein is kept open and an active fixation RV lead, with a straight stylet in position, is advanced down to the right ventricular apex under fluoroscopy guidance. Due caution is taken when advancing the lead and lead advancement is stopped if there is any resistance.
5. As the lead crosses the tricuspid valve and is advanced into the RV, the ECG is monitored for the presence of ventricular ectopics suggestive of crossing the valve into the RV.

△ Advancement of the lead is performed cautiously across the tricuspid valve. If the lead does not directly cross the valve, the lead is prolapsed with the stylet retracted ∼5cm - 8cm followed by straightening out the lead with the stylet fully inserted.

6. Once at the RV apex, the active fixation lead is deployed by applying clockwise turns on the proximal end of the lead to screw in the lead.
7. With the analyser connected to the distal end of the lead, the lead parameters are tested in the bipolar configuration. Target parameters include an R wave > 6mV, an impedance value between 300–1500Ω and a pacing threshold of <1V.
8. The lead is secured with 2.0 silk ties at the proximal and distal lead cuffs. The distal cuff is secured with the jugular vein simultaneously achieving vein closure and haemostasis.
9. The lead is connected to the appropriate port of the generator.
10. The subcutaneous pocket for the generator is created. The site needs to be sufficiently distal to the original incision towards the shoulder. The generator with the residual lead coiled is placed into the pocket.
11. The pocket is closed with interrupted 2.0 Vicryl sutures.
12. The VT, VF and FVT zones are programmed for detection only and all therapy is turned off. The rationale behind this is to avoid inappropriate shocks as the higher sinus rates and T wave oversensing in the model.
13. The device is then left connected wirelessly to the programmer in the emergency mode to allow prompt defibrillation of intra-operative life-threatening ventricular arrhythmias.

#### C. Induction of myocardial infarction

△ All the MI induction equipment (except the indeflator and the balloon) listed above should be pre-flushed with heparinised saline solution (i.e., 500mL sodium chloride 0.9% solution containing 10,000 IU heparin prepared in a sterile kidney dish) and prepared as follows:

• Insert the dilator into the 6 Fr haemostatic introducer sheath.
• Connect the 6F JR4 guide catheter to the Honor® Hemostasis Valve, which is connected to a 3-way valve, and then insert the 0.35 J tipped guide wire through the Hemostasis Valve all the way to the end of the catheter.
1. The previously identified carotid artery is the vascular access site for this part of the procedure.
2. The proximal suture is tied off.
3. The vessel is controlled with the previously placed proximal and distal ties.
4. A 14G cannula is advanced into the carotid artery towards the distal end.

△ The cannula is cautiously inserted to remain intraluminal and avoid transecting the carotid artery.

5. Once intravascular access is achieved, the x short guide wire is advanced down the cannula and exchanged for the 6F 12cm haemostatic sheath using the Seldinger technique.
6. The sheath is loosely secured via the side loop to prevent displacement secondary to the carotid pulsation.
7. Upon achieving arterial access, a 10,000 IU bolus if IV heparin is administered followed by a maintenance infusion of 10IU/ml to reduce the risk of thrombotic complications with the indwelling arterial equipment.
8. 30 minutes prior to coronary access, amiodarone 100mg IV is administered followed by a maintenance bolus dose of 50mg/hour.
9. Via the introducer sheath, a 6F Guide Judkins Right (JR4) catheter is advanced with a pre-loaded wire 0.35 in 150cm J wire. This is introduced under fluoroscopic guidance with the J wire leading to reduce vascular trauma.
10. The J wire is advanced down to the aortic valve. When it reaches the aortic valve, mild resistance should be felt, which corresponds to the J wire looping at the valve level.
11. Then the guide catheter is advanced over the wire down to the aortic valve level and the wire is removed.
12. A 50ml syringe with 50% contrast mix is connected to the end of the guide catheter.
13. Using fluoroscopic guidance, the catheter is advanced into the left coronary system with catheter motion and contrast injection used to confirm/identify position.

△ Engagement of left coronary artery system is done gently under direct fluoroscopic guidance observing catheter tip motion and a gentle contrast injection to ensure the ostium of the vessel is not dissected and to avoid deep intubation of the coronary.

14. Once engaged adequately, coronary angiography is performed with the 50% contrast mixture to delineate the left coronary anatomy identifying the LAD coronary artery and the second diagonal branch (D2). This will guide identification of the occlusion target which is in the LAD after the D2 branch.
15. A bolus dose of 100mg Lidocaine is administered intravenously 20-30 minutes prior to coronary occlusion via the peripheral cannula.
16. Coronary engagement is maintained with the guide catheter, whilst a 0.0014 in wire (normal length wire) is advanced down to the distal LAD. The wire is introduced into the haemostasis valve via the introducer needle provided in the set.

△ The wire is advanced gently under fluoroscopic guidance to avoid coronary perforation. Avoid buckling the wire tip as it is advanced.

17. The indeflator is prepared with the 50:50 contrast and heparinised saline solution mix. For this, the chamber of the indeflator is filled with the mixture by aspiration, then the handle is turned clockwise to expulse the fluid, removing any leftover bubbles and leaving the mixture bubble-free.
18. Depending on the approximated diameter of the vessel (which is determined by comparing the vessel calibre to the guide catheter, which represents a width of approximately 2 mm), an appropriate size intracoronary balloon is selected.
19. The balloon’s chamber is filled with contrast to ensure there is no air and is connected to the indeflator. A vacuum is created by adding negative pressure and holding it in place to empty the balloon and ensuring that there is no air within. The vacuum must be maintained during the balloon’s introduction.

△ Ensure that the device is undamaged.

Do not pre-inflate or test the balloon before insertion.

20. The protective sleeve is removed from the balloon. The balloon catheter has a central lumen, which allows it to be advanced over the 0.0014 in wire.
21. The 0.0014 in wire is fixed in the distal LAD and the intracoronary balloon is advanced to the target site of occlusion over this wire whilst maintaining the distal position of this wire at all times.

△ As the balloon is advanced, it is critical to hold in place the thin wire, so it does not cause coronary trauma or perforation by inadvertent advancement.

22. The indeflator is used to inflate the intracoronary balloon at the target occlusion point within the LAD (i.e., immediately after the D2 vessel bifurcates). The contrast mixture is filled into the intracoronary balloon by the indeflator, resulting in the occlusion.

△ Inflation of the intracoronary balloon is performed slowly to ensure adequate but not excessive inflation which can cause coronary artery damage.

23. A coronary angiogram is performed to confirm that there is no visible flow of contrast distal to the occlusion point suggesting adequate balloon inflation and this inflation is maintained for a 90-minute duration.
24. A further bolus intravenous dose of 50mg Lidocaine is administered via the peripheral cannula at 20 minutes post occlusion.

△ Continuous monitoring of ECG, blood pressure, and oxygen saturations are necessary during this time.

25. Typically, animals begin to show ST changes almost immediately after the occlusion, with ventricular arrhythmias happening between 20-40 minutes later. Any ventricular arrhythmias need to be quickly treated by internal cardiac defibrillation using the 35J cardiac defibrillator device that has been implanted in the beginning of the procedure. This is done manually, using the header of the ICD programmer, since the device is in emergency mode.

△ Prompt defibrillation is necessary upon recognition of ventricular arrhythmia. This may sometimes require multiple defibrillations. Therefore it is important to ensure that the implanted device has sufficient battery life.

26. Coronary perfusion is restored by deflation of the balloon at 90 minutes with a coronary angiogram confirming re-perfusion down the coronary artery.
27. Atenolol 1mg is administered intravenously via the peripheral cannula on reperfusion.
28. The guide, balloon and wire are removed from the heart.
29. The carotid sheath is removed and the carotid artery is tied off with a 2.0 silk suture achieving haemostasis.

△ The removal of the carotid sheath is performed simultanously as the carotid is tied off to avoid excessive bleeding.

30. The wound is closed in layers with a 2.0 monocryl absorbable suture.
31. The animal is then gradually awakened, extubated, and recovered.
32. The animal is checked one hour after recovering from the procedure. Observations are made from a safe distance to confirm that it is still alive, aware, and moving around in the pen.
33. On the first day, any interaction with the animal is limited to prevent frightening it and arrhythmias.

