## Supplemental Figure 1 for "A novel minimally invasive and reproducible large animal ischaemia-reperfusion-infarction model: methodology and model validation"

### Slide 1
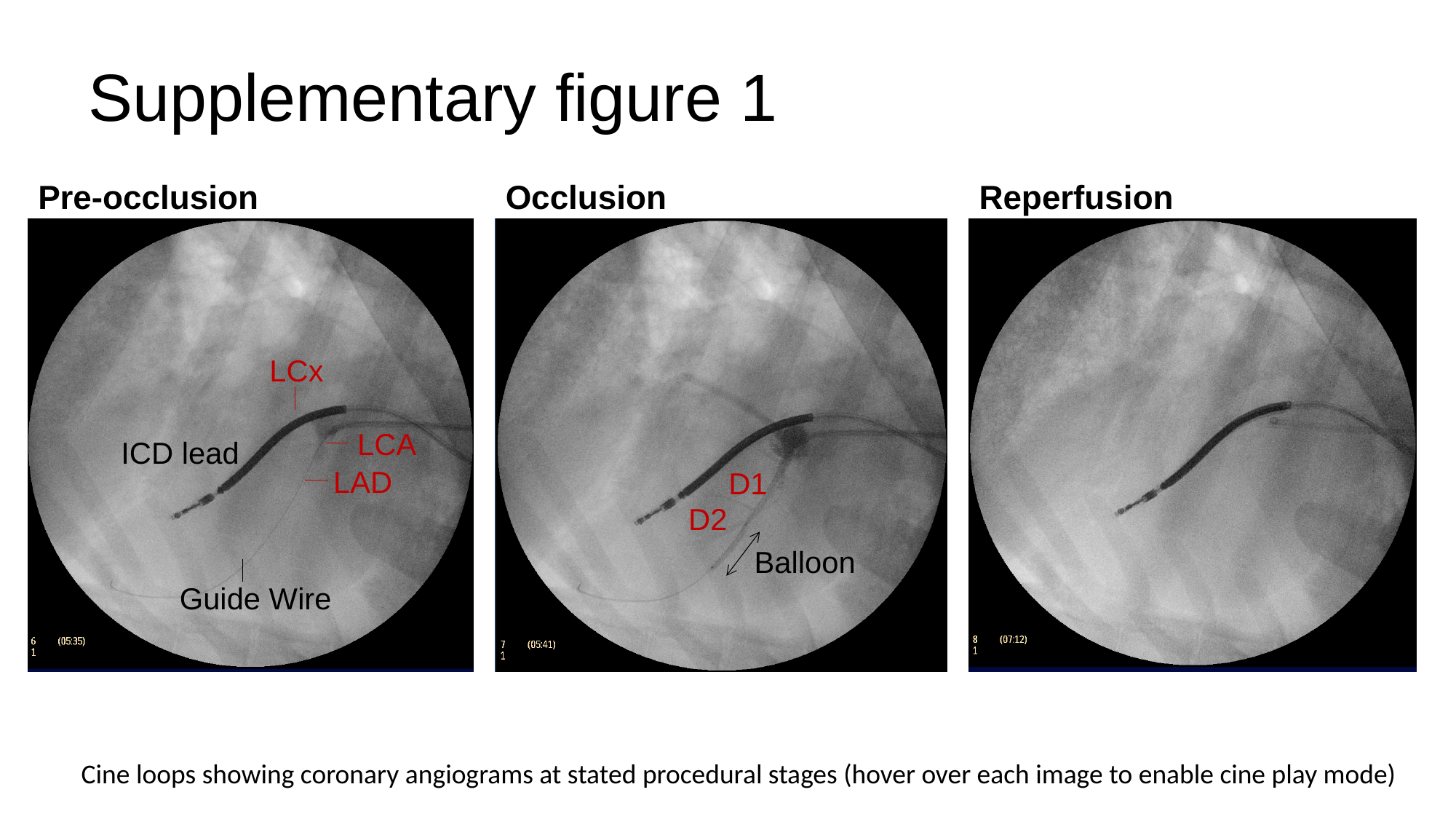

#### Supplementary figure 1
Pre-occlusion
LCx
LCA
ICD lead
LAD
Guide Wire
Occlusion
Reperfusion
Weeks
D1
D2
Balloon
Cine loops showing coronary angiograms at stated procedural stages (hover over each image to enable cine play mode)
