## Supplemental Table 1 for "A novel minimally invasive and reproducible large animal ischaemia-reperfusion-infarction model: methodology and model validation"

| | Alb (g/L) | TP (g/L) | Gluc (mmol/L) | ALP (U/L) | AMY (U/L) | BUN (mmol/L) | Crea (µmol/L) | Ca (mmol/L) | Phos (mmol/L) | Na (mmol/L) | K (mmol/L) | Glob (g/L) | Urea (mg/dL) | Alb/Glob ratio | BUN Crea ratio | NaK ratio |
|---|---|---|---|---|---|---|---|---|---|---|---|---|---|---|---|---|
| **baseline** | 33.71 ± 0.61 | 72.14 ± 2.11 | 6.23 ± 0.69 | 162.43 ± 11.42 | 23.67 ± 5.3 | 6.5 ± 0.67 | 102.14 ± 5.14 | 2.4 ± 0.08 | 2.11 ± 0.21 | 145.57 ± 1.39 | 5.47 ± 0.16 | 38.43 ± 2.34 | 6.5 ± 0.67 | 0.9 ± 0.07 | 16.17 ± 2.11 | 26.57 ± 0.65 |
| **90 mins** | 31 ± 1.13 § | 62.14 ± 1.9 | 5.06 ± 0.89 | 193.29 ± 34.31 | 25.86 ± 5.16 | 7 ± 0.31 | 112.43 ± 6.12 | 2.03 ± 0.12 § | 2.12 ± 0.2 | 147.14 ± 1.62 | 4.99 ± 0.44 | 31.14 ± 1.45 $ | 7 ± 0.31 | 1.01 ± 0.06 | 15.69 ± 1.08 | 30.43 ± 2.17 |
| **day 3** | 34.57 ± 0.81 | 72.29 ± 1.87 | 4.99 ± 0.37 | 133 ± 21.01 | 28.29 ± 4.41 | 5.34 ± 0.3 | 92.14 ± 4.19 | 2.55 ± 0.09 | 1.93 ± 0.1 | 151 ± 1.2 | 5.66 ± 0.23 | 37.71 ± 1.57 | 5.34 ± 0.3 | 0.91 ± 0.03 | 14.56 ± 1.05 | 27 ± 1.25 |
| **week 1** | 32.14 ± 1.35 | 72.57 ± 1.46 | 8 ± 1.03 | 139.29 ± 11.65 | 28 ± 4.27 | 5.83 ± 0.57 | 97 ± 3.93 | 2.55 ± 0.05 | 2.11 ± 0.17 | 147.43 ± 1.65 | 5.3 ± 0.12 | 39 ± 1.31 | 5.83 ± 0.57 | 0.87 ± 0.03 | 15.01 ± 1.6 | 27.71 ± 0.68 |
| **week 3** | 34.57 ± 0.9 | 72.86 ± 1.44 | 6.83 ± 0.41 | 170.14 ± 21.58 | 27.43 ± 4.1 | 6.49 ± 0.64 | 89.57 ± 5.18 | 2.58 ± 0.06 | 2.03 ± 0.11 | 150.14 ± 0.7 | 5.2 ± 0.1 | 38.29 ± 1.08 | 6.49 ± 0.64 | 0.93 ± 0.03 | 18.27 ± 1.87 | 28.86 ± 0.55 |
| **week 8** | 36.43 ± 0.75 | 71.29 ± 1.98 | 5.87 ± 0.76 | 177.57 ± 15.02 | 30 ± 3.17 | 5.9 ± 0.6 | 82.14 ± 3.61 * | 2.58 ± 0.05 | 1.99 ± 0.12 | 152.14 ± 1.62 * | 5.24 ± 0.18 | 34.86 ± 1.87 | 5.9 ± 0.6 | 1.06 ± 0.06 * | 17.71 ± 1.58 | 29.14 ± 0.99 |

**Supplementary Table 1. Summary of procedural changes in plasma biochemistry profile.**
***, $p < 0.05$; $, $p < 0.001$ and §, $p < 0.0001$ versus baseline**
